# Exogenous Sphingomyelinase Mediates MSC-derived EV Biogenesis and Enhances Potency via Repackaging of Molecular Cargo

**DOI:** 10.1101/2025.08.21.671536

**Authors:** Daniel C. Shah, S’Dravious A. DeVeaux, Hongmanlin Zhang, Alan Y. Liu, Tosin A. Adedipe, Nathan F. Chiappa, Afra I. Toma, Krishna A. Patel, Young C. Jang, Steven L. Goudy, Todd Sulchek, Krishnendu Roy, Edward A. Botchwey

## Abstract

Mesenchymal stromal cells (MSCs) exert regenerative and immunomodulatory effects largely through secreted paracrine factors and extracellular vesicles (EVs), which transfer proteins, lipids, and nucleic acids to recipient cells. Lipid composition critically influences EV stability, uptake, and bioactivity. Sphingomyelinase (SMase), an enzyme that hydrolyzes sphingomyelin into ceramide, regulates EV biogenesis by inducing membrane curvature and initiating inward membrane budding. Here, MSCs were treated with SMase, and EVs were isolated and characterized by nanoparticle tracking analysis, miRNA sequencing, lipidomics, and proteomics. SMase treatment increased EV yield and altered lipid, protein, and miRNA cargo linked to TNF-α signaling, wound healing, and angiogenesis. Functionally, SMase-EVs suppressed TNF-α in macrophages, showed trending increased HUVEC tubular formation, and altered T-cell populations following local delivery in a critical murine oral wound defect model. These findings highlight how enzymatic lipid remodeling modifies MSC-EVs, enhancing their therapeutic potential and informing strategies for optimized EV-based therapies and scalable production.

## INTRODUCTION

Mesenchymal stromal cells (MSCs) have emerged as a compelling cell therapy candidate due to their robust immunomodulatory and pro-regenerative properties, particularly in the context of chronic inflammatory, autoimmune, and degenerative diseases [1]. They can be isolated from multiple tissue sources such as bone marrow, adipose tissue, umbilical cord, and dental pulp; each offering distinctive biological advantages related to angiogenesis, immune regulation, extracellular matrix (ECM) remodeling, and cell proliferation [2, 3]. Despite initial emphasis on their multi-lineage differentiation capabilities, poor engraftment, retention, and survival of MSCs *in vivo* have significantly hampered their therapeutic impact [4, 5]. Furthermore, limited direct evidence of *in vivo* differentiation challenges the notion of MSC-based tissue replacement as a primary mechanism of action [6, 7]. Additionally, ex vivo culture variability, donor heterogeneity, and the lack of well-defined critical quality attributes have further constrained the clinical efficacy of MSCs [8].

Increasingly, research points to MSCs’ paracrine effects, rather than long-term engraftment, as the main driver of their therapeutic properties. MSCs secrete a diverse repertoire of cytokines, growth factors, lipid mediators, mRNA, microRNA (miRNA), and extracellular vesicles (EVs), all of which orchestrate immunomodulation and promote tissue repair [9, 10]. This paracrine focus has led to a paradigm shift toward harnessing the MSC secretome, particularly small EVs, as a more feasible, cell-free therapeutic alternative with fewer concerns related to tumorigenicity, immune rejection, and large-scale manufacturing [11]. Small EVs, often referred to as exosomes (30–150 nm), are enriched in proteins, nucleic acids, and lipids, which confer potent immunomodulatory and pro-regenerative functions. As of mid-2025, over 30 clinical trials registered on clinicaltrials.gov have investigated MSC-derived EVs/exosomes in treating neurodegenerative disorders, inflammatory diseases, diabetes, and kidney and skin conditions [12, 13].

The potency of EVs is intimately tied to their lipid and protein composition, as well as to the organelle-level dynamics that govern EV formation. EV biogenesis can proceed through the endosomal sorting complex required for transport (ESCRT)-dependent or ESCRT-independent pathways, both of which rely on localized membrane remodeling to generate intraluminal vesicles (ILVs) that ultimately fuse with the plasma membrane (PM) to be released as exosomes [14]. During this process, lipids such as ceramide, phosphatidylserine, and cholesterol can help create curvature and assemble specialized membrane microdomains involved in cargo sorting known as lipid rafts [15–18]. These microdomains provide a platform for organizing proteins and lipids, facilitating the selective enrichment of cargo into EVs often through glycosylphosphatidylinositol (GPI)-anchor sites [19, 20]. The role of lipid rafts in ensuring selective cargo incorporation underscores their significance in enhancing the therapeutic potential of EVs. This interplay between membrane architecture, lipid raft formation, and the ESCRT-independent ceramide pathway is increasingly recognized as a pivotal regulatory axis for EV formation [21–24].

Among the sphingolipids, ceramide (Cer) plays a central role in ESCRT-independent exosome formation by modulating membrane curvature within multivesicular bodies (MVBs) [25, 26]. Sphingomyelinase (SMase) catalyzes the hydrolysis of sphingomyelin (SM) to produce Cer, thereby promoting inward budding of the MVB membrane and the eventual release of exosomes [27]. Endogenous sphingolipid metabolism is crucial not only for membrane structure but also for their potency and selective incorporation of bioactive cargo, including microRNAs (miRNAs) and proteins into EVs [15, 24, 28–30]. Although endogenous SMase activity is known to be essential for intraluminal vesicle budding, the therapeutic exploitation of exogenous SMase to enhance Cer generation and improve EV output remains largely unexplored. Studies in other cell types have demonstrated that exogenous Cer or Cer modulators (e.g., C6-ceramide) can reshape EV cargo composition to alter angiogenic and immunomodulatory properties [22, 31, 32]. In addition, endogenous lipid raft formation and SMase activity has been shown to have an intrinsic relationship that can directly contribute to alterations in cellular biophysical properties [33]. However, it is yet to be determined whether exogenous SMase can similarly restructure MSC membrane domains, selectively augment certain subpopulations of EVs, and confer superior immunomodulatory or pro-angiogenic activity.

Although MSC-derived EVs have garnered attention for their safety and therapeutic potential, key questions remain regarding the impact of exogenous SMase-induced membrane remodeling on EV biogenesis and cargo sorting, the specific lipid raft reorganization that enhance EV cargo sorting, and the feasibility of harnessing these modifications to strengthen angiogenic and immunosuppressive outcomes. Our previous findings have demonstrated that exogenous SMase treatment of MSCs reprograms cellular sphingolipid metabolism without inducing atypical levels of senescence, autophagy, or apoptosis leading to promising insights on maintaining cellular viability [34]. These findings indicate that SMase not only acts as a lipid metabolic regulator but also primes MSCs by augmenting secretory pathways linked to EV biogenesis and immunoregulatory factor release. We hypothesize that SMase treatment reconfigures sphingolipid metabolism and lipid raft organization, preferentially activating ceramide-dependent EV biogenesis and increasing lipid microdomain sites for binding of trafficking proteins, allowing for packaging of immunomodulatory proteins and miRNAs. To investigate these mechanisms, we employed a multi-omic characterization to include lipidomics, proteomics, and miRNA transcriptomics combined with high-resolution imaging of membrane architecture and EV potency assessment. This systematic approach illuminates how sphingolipid perturbations may alternatively govern EV assembly and cargo selection, laying the groundwork for refining MSC- or EV-based therapeutics and informing scalable manufacturing processes.

## MATERIALS AND METHODS

### MSC expansion and culture

BM-MSC donor RB183 was purchased from RoosterBio, Inc. (Frederick, MD) with a population doubling level (PDL) of 8.9. These cells were considered as passage 0 (P0). The MSCs were thawed and expanded by protocols provided by RoosterBio, Inc. Briefly, cells were cultured in RoosterNourish^TM^ media (High Performance Media Kit, KT-001) containing serum-derived RoosterBooster^TM^ in T-225 flasks at an approximate density of 3,333 cells/cm^2^. Cells were grown to 80% confluence, harvested and cryopreserved per manufacturer’s protocol for sub-passaging or downstream assays. Once sub-passaged cells were ∼80% confluent, they were treated with or without 0.3 U/mL of filter sterilized sphingomyelinase (S7651, Sigma-Aldrich, St. Louis, MO). This phospholipase enzyme was produced by *Bacillus cereus* and belongs to the neutral sphingomyelinase enzyme family. After approximately 24 hrs, the treatment was removed, and the cells were washed with PBS. MSCs for all experiments were used at P1-P3 and a PDL of ∼12-18.

### Atomic Force Microscopy

The cell stiffness measurements were performed with an MFP-3D Bio-AFM (Asylum Research). MSCs were seeded at 10,000 cells/cm^2^ in a Fluorodish (PI) and allowed to attach overnight prior to SMase treatment that was applied as previously described. A 5.24 µm spherical silica bead was attached to a tipless silicon nitride cantilever with calibrated stiffness of 0.0204 N/m (MLCT-C from Bruker, nominal spring constant k = 0.01 N/m) using a two-part epoxy and cured for 48 hrs. The thermal method was used to calibrate the cantilever spring constant immediately prior to use by indenting the glass bottom of a Fluorodish and performing a Lorentzian fit to the thermal spectrum [35]. The cantilever probe was visually aligned with the MSCs center using an integrated inverted optical microscope and translated with a vertical velocity of 4 um/s to indent the cell with increasing force until a 5 nN trigger force was reached. One measurement per cell was obtained, 19 untreated MSCs and 21 SMase treated MSCs were probed, respectively. A Hertzian contact model was used to calculate Young’s modulus for stiffness characterization through customized PYTHON code (www.github.com/nstone8/pyrtz). The early contact portion (10% of the force trigger) was used to calculate cell stiffness.

### Fluorescent Tracer Assay of Secreted Lipid Metabolites and Exchange Flux Modeling

As previously described, MSCs were cultured and treated with SMase for 24 hrs before incubated with 10 µM of C6 NBD-sphingomyelin (810218, Avanti Polar Lipids) for 2 hrs [36]. Conditioned media was collected for lipid extraction and resuspended in mobile phase (850:150:15 MeOH:H_2_O:H_2_PO_4_) and analyzed using a Prominence High Performance Liquid Chromatography (HPLC) (Shimadzu UFLC, flow rate at 1mL min-1, excitation = 460 nm, emission = 535 nm) equipped with a C18 250 x 4.6 mm LC column (Phenomenex, Torrance, CA). Lipid concentrations were normalized by total protein using a Pierce™ BCA Protein Assay Kit (23225, Thermo Scientific™) according to the manufacturer’s protocol. Dynamic flux estimation was conducted in MATLAB 2023b (Mathworks) as previously described [36].

### Extracellular vesicle isolation

Bone Marrow-derived MSCs were cultured as previously described. Once at ∼80% confluency, the flasks were replaced with 30 mLs of RoosterCollect^TM^-EV (M2001, Rooster Bio.) with or without 0.3 U/mL of SMase. Conditioned media was harvested after 24 hrs and immediately centrifuged at 300×g for 10 mins to remove cells. The supernatant was subsequently spun down at 2000×g for 30 mins to remove dead cells. The resulting conditioned media was then filtered through a 0.22 µm filter and stored at −80 °C. Once needed, the samples were thawed on ice overnight at 4 °C. The samples were transferred to ultracentrifuge tubes (361625, Beckman Coulter) and spun at 10,000×g for 30 mins using a SW 32 TI rotor (Beckman Coulter) to remove cellular debris and large vesicles. The supernatant was transferred to a new ultracentrifuge tube and was centrifuged at 100,000×g for 90 mins. The resulting pellet was washed with cold PBS and spun down at 100,000×g for 90 mins before being resuspended in 100 µl of 0.10 µm filtered cold PBS for downstream characterization and assays.

### Nanoparticle Tracking Analysis

To measure their size profile and concentration, isolated EVs were diluted 1:100 in 0.10 µm filtered PBS and Nanoparticle Tracking Analysis (NTA) was performed using the Nanosight NS300 (Malvern Instruments). Five 60 sec videos were captured with the syringe pump set to 100, camera level set to 13, and screen gain set to 7. Data was analyzed with NTA 3.4 Build 3.4.4 software with detection threshold set to 6. A sample of diluent was also measured and used for background subtraction in downstream data processing.

### Transmission Electron Microscopy

To visualize the morphology of the EVs, 10 µl of sample was placed on a 400-mesh carbon-coated copper grid (Electron Microscopy Services) for 10 min. The remaining volume was blotted away using filter paper. The grid was stained with 1% aqueous uranyl acetate for 30 sec before being blotted away. Immediately, the grids were rinsed with 10 µl of deionized water and quickly removed. The grid was allowed to dry prior to being placed in a Hitachi HT7700 120kV TEM (Hitachi High-Technologies Corporation).

### Zeta Potential

EVs were diluted in 0.1 µm filtered PBS 1:100 and placed into a folded capillary zeta cell (Malvern Panalytical). The zeta cells were placed into a Zetasizer Nano Z (Malvern Instruments) to measure the zeta potential of each sample with the system settings set to 50 zeta runs.

### Flow cytometry of classical EV markers

EVs were prepared as previously discussed and diluted 1:1000 in 0.1 µm filtered PBS. EVs were labeled with CellTrace^TM^ Blue (C34568, Thermo Fisher Scientific), CD63 Brilliant Violet 421 (353029, BioLegend®), CD9 Alexa Flour 647 (MA5-18154, Thermo Fisher Scientific), and CD81 PE (349506, BioLegend®); all being diluted 1:1000 and incubated for 10 mins at RT then quickly put on ice. Samples were run on a Cytek Aroura (Cytek Biosciences) with all default voltages increased by 100% [37–39].

### Immuno-Staining and Confocal Microscopy

MSCs were seeded onto µ-slides (80826, ibidi) at 10,000 cells/cm^2^ and allowed to attach overnight. Then were treated with SMase as previously described. Following this treatment, MSCs were stained to identify lipid membrane alterations with Vybrant™ Alexa Fluor™ 555 Lipid Raft Labeling Kit (V-33404, Invitrogen™) as per the manufacturer’s instructions and fixed with 4% PFA for 15 min at RT. The cells were then incubated in a 2% BSA, 0.5% goat serum, 0.5% Triton-X 100 blocking solution for 30 min at RT followed by an overnight incubation at 4 °C of CD63 Alexa Fluor™ 488 (MA5-18149, Invitrogen™) diluted 1:100 in blocking solution. 0.1% Tween was used to wash the cells three times, and CellMask™ Deep Red Actin Tracking Stain (A57245, Invitrogen™) was diluted 1:1000 and incubated for 15 min at RT. Hoechst 33342 (H1339, Invitrogen™) was diluted 1:2000 and incubated for 10 min at RT to visualize the nuclei.

Z-stack fluorescent images were acquired using a Nikon W1 Spinning Disk Confocal microscope using a 60X oil objective (1.4 numerical aperture, working distance of 0.13 mm). Five random fields of view (253.44 μm × 253.44 μm) were taken from each well for image analysis in Fiji. For lipid raft and CD63 expression quantifications, thresholds were applied to identify the fluorescent signal area, which was measured and normalized to the cell number in each image. Data from the five images per well were averaged to generate representative values for each well (n=4).

### Lipidomic LC-MS profiling of MSCs and EVs

MSCs and EVs (n=3) were stored in −80 °C before being thawed on ice. MSCs and EVs were transferred to 1.5 mL Eppendorf tubes containing 100% isopropanol. Glass beads (0.5 micron) were added to the tubes and the cells and EVs were lysed using a Qiagen TissueLyser II for 10 min at 30 Hz. The samples were dried by vacuum centrifugation and reconstituted with isopropanol containing isotopically labelled internal standards (Table S1) at a ratio of 40 µL/10^9^ EVs and 40 µL/10^6^ cells. Lipids were analyzed with an ultraperformance liquid chromatography system using a Vanquish Horizon (Thermo Fisher Scientific) equipped with a CORTECS Premier T3 column (Waters Corp, 2.1×100 mm, 1.6µm) coupled to an Orbitrap Exploris 240 mass spectrometer (Thermo Fisher Scientific). The mobile phases were water with 10mM ammonium acetate and 0.05% acetic acid (mobile phase A) and 20:80 acetonitrile:isopropyl alcohol with 10 mM ammonium acetate and 0.05% acetic acid (mobile phase B). The chromatographic method used the following gradient program: 0 mins 70% A; 0.5 mins 70% A; 1.4 mins 40% A; 7.5 mins 0% A; 10.2 mins 0% A; 10.3 mins 70% A, 12 mins 70% A. The flow rate was set at 0.30 ml/min. The column temperature was set to 45 °C, and the injection volume was 2 µL. Full scan data was acquired in positive mode with a filter of 150-2000 m/z at 120,000 resolution. MS/MS data was collected in data-dependent acquisition (DDA) with an isolation window of 0.8 m/z, stepped HCD collision energies of 15, 30, 50%, and an orbitrap resolution of 15,000. Compound Discoverer V3.3 (Thermo Fisher Scientific) was used to process raw LC-MS data. Data processing steps included peak detection, spectral alignment, grouping of isotopic peaks and adduct ions, drift correction, and gap filling. Lipid annotations were accomplished based on accurate mass and MS2 fragmentation pattern matching to local spectral databases built from curated experimental data and/or accurate mass matching to Lipid Maps. Differentially expressed annotated lipids were analyzed using lipid pathway enrichment analysis [40].

### Proteomic LC-MS profiling of MSCs and EVs

To extract the proteins, the cell pellet and extracellular vesicle pellet was resuspended in 200 µl of SDS lysis buffer (5% SDS, 50 mM TEAB pH 8.5) supplemented with Pierce Protease and Phosphatase Inhibitor Mini Tablets, EDTA-free (A32961, Thermo Scientific™) and incubated at 95 °C for 15 min. The tube was spun at 10,000×g for 5 min and the supernatant was transferred into a new tube. The supernatant was further clarified by centrifuging it at 21,000×g for 10 min at 4 °C. The protein extract was removed from the top into a fresh tube and protein concentration was determined by BCA method as per the manufacturer’s instructions. An aliquot of 1 µg/µL in 50 µL solutions of proteins was used for proteolytic digestion. Micro s-Trap (Protifi) columns were used for tryptic digestion as per the manufacturer’s instructions. In brief 50 µg of dried protein was resuspended in 23 µl of SDS lysis buffer. Proteins were reduced by adding TCEP (final concentration 5 mM TCEP) and incubating them at 55 °C for 15 min. Reduced samples were alkylated by adding Iodoacetamide at a final concentration of 10 mM. Samples were acidified by adding phosphoric acid (final concentration ∼2.5%). 165 µl of binding buffer (100 mM TEAB (final) in 90% methanol) were added to each tube and the total contents were transferred to s-Trap columns. These tubes were spun at 4,000×g for 1 min and the flow through was discarded. Trapped proteins were washed three times with wash buffer (100 mM TEAB (final) in 90% methanol). Sequencing grade trypsin (Thermo Scientific) were added to the trap in a 1:20 trypsin to protein ratio in 30 µl of 100 mM TEAB. The S-trap columns were incubated at 37 °C overnight. On the following day, digested peptides were eluted sequentially by 50 mM TEAB, 0.1% Formic acid and 50% Acetonitrile. Pooled eluates were dried in speedvac and cell lysate samples were reconstituted in 25 µl of 0.1% FA solution. EV peptide samples were reconstituted in 100 mM TEAB and labeled with isobaric mass tags using the TMTpro 16 plex label reagent set (A44521, Thermo Scientific) per the manufacture’s protocol. An externally calibrated Thermo Q Exactive Plus (high-resolution electrospray tandem mass spectrometer) was used in conjunction with Dionex UltiMate3000 RSLCnano System. A 2 μL (1 μg equivalent) sample was aspirated into a 50 μl loop and loaded onto the trap column (Thermo µ-Precolumn 5 mm, with nanoViper tubing 30 µM i.d. × 10 cm). The flow rate was set to 300 nL/min for separation on the analytical column (Acclaim pepmap RSLC 75 μM × 15 cm nanoviper). Mobile phase A was composed of 99.9% H_2_O (EMD Omni Solvent), and 0.1% formic acid and mobile phase B was composed of 99.9% ACN, and 0.1% formic acid. A 120 min linear gradient from 3% to 45% B was performed. The LC eluent was directly nanosprayed into Q Exactive Plus mass spectrometer (Thermo Scientific). During the chromatographic separation, the Q Exactive plus was operated in a data-dependent mode and under direct control of the Thermo Excalibur 3.1.66 (Thermo Scientific). The MS data were acquired using the following parameters: 20 data-dependent collisional-induced-dissociation (CID) MS/MS scans per full scan (400 to 1800 m/z) at 70,000 resolutions. MS2 was acquired at 17,500 resolutions. Ions with single charge or charges more than 7 as well as unassigned charge were excluded. A 10 sec dynamic exclusion window was used. All measurements were performed at room temperature. Resultant raw files were searched with Proteome Discoverer 3.0 using Sequest™ HT as the search engine using species specific FASTA database and percolator as peptide validator. STRING version 12 and Cytoscape version 3.10.2 were used to construct the protein-to-protein interaction maps and gene ontology (GO) enrichment analysis.

### miRNA enriched RNA-sequencing

A QIAcube (Qiagen) automatic sample handler was used with using the miRNeasy Tissue/Cell Advanced Mini Kit (217604, Qiagen) to extract total RNA from each of the cell and EV pellets (n=3 per group). Quantity and quality of RNA was checked using Agilent RNA 6000 pico kit (5067-1513, Agilent) on the Agilent 2100 Bioanalyzer (Agilent Technologies). Following the manufacturer’s protocol, sequencing libraries were prepared using the QIAseq miRNA library kit (331502, Qiagen). Equivalent RNA concentrations were made and sequenced with a NextSeq 550 (Illumina) in mid output mode, single-end reads were made with a read length of 72 bp. Raw sequencing data was processed using Qiagen’s online RNA portal (https://rnaportal.qiagen.com). The samples were aligned and mapped to the homo sapiens reference miRbase_v22 using the RNA-seq Analysis Portal 5.0. Volcano plots and heatmaps were constructed using FDR ≤ 0.1 and |Log_2_ fold change| > 2. Differentially expressed miRNAs (DEmiRs) were then used to identify mRNA targets via mirecords, mirtarbase, and tarbase using the multiMiR package in R version 4.3.3. KEGG enrichment pathways were then mapped using the ClusterProfiler package and chord plots were generated using the GOplot package.

### Macrophage Activation Assay

RAW 264.7 (TIB-71, ATCC) murine macrophages were seeded at 40,000 cells/cm^2^ and cultured in RPMI 1640 medium (11875093, Gibco^TM^) supplemented with 10% heat inactivated FBS and 1% penicillin-streptomycin. Once ∼80% confluent, cells were treated with or without 50 ng/mL of lipopolysaccharide and co-incubated with EVs for ∼24 hrs using approximately 1e^9^/mLfrom either UT or SMase derived conditions (n=3). Conditioned media (CM) was collected and immediately stored at −80°C for downstream analysis. To quantify TNF-α secretion, CM was thawed on ice and assayed using a Mouse TNF alpha ELISA Kit (BMS607-3, Invitrogen™) according to the manufacturer’s protocol. Samples were diluted at 1:5 and measured using a microplate reader at 450 nm, readings were background subtracted and fit to the standard curve.

### HUVEC Angiogenesis Assay

Lonza’s pooled human umbilical vein endothelial cells (HUVECs) (NC1835181, Fischer Scientific) were expanded using EGM-2 MV Endothelial Cell Growth Medium-2 Bullet kit with SingleQuots supplement pack (NC9902887, Fischer Scientific) for 2 passages. To obtain conditioned media, MSCs were grown in T-225 using RoosterNourish^TM^ until 80-90% confluency. Media was then aspirated, washed with PBS, and fresh EGM-2 MV media was added with the appropriate treatment for 24 hrs. The conditioned media was spun down at 500×g for 5 min before the supernatant was removed and frozen until use. The EVs were isolated as previously described. 75 µL of Matrigel (#354234, Corning) was added to a flat bottom 96-well plate and placed in incubator for 1.5 hrs to allow for polymerization. HUVECs were then harvested, resuspended in conditioned or EV containing media and seeded at 28,800 cells/cm^2^. 50 µg of EVs were used after measured with Micro BCA™ Protein Assay Kit (23235, Thermo Scientific™). After 24 hrs, HUVECs were imaged using a 4x phase contrast microscope 6-7 images were taken per well and the parameters were averaged. ImageJ angiogenesis plug in was used to analyze images.

### Synthetic Hydrogel Synthesis

PEG-based synthetic hydrogels were synthesized using 4% (w/v) maleimide end-functionalized (four-arm) PEG macromer (PEG-4MAL) (4ARM-PEG-MAL-20K, Laysan Bio), functionalized with a thiol-containing adhesive peptide GCGYGRGDSPG (RGD) (RP20297, GenScript), crosslinked with the protease-degradable peptide VPM (GCRDVPMSMRGGDRCG) (AAPPTec) in 25 mM HEPES (pH ∼5.5). RGD and VPM were used at a final concentration of 1.0 mM and 3.3 mM, respectively. Hydrogels were loaded with approximately 7e^8^ EVs. All chemical components were sterile filtered through a spin column following pH measurements and kept in aseptic conditions prior to local palate administration.

### Critical Murine Oral Wound Defect Model

In accordance with Georgia Institute of Technology’s Institutional Animal Care and Use Committee (approval no. BOTCHWEY-A100669-04, date: 04/18/2024), C57BL/6 mice (Jax Lab, age 4-6 weeks) (n = 3-4/group) were allowed to acclimate for 1-week and housed under precise conditions: temperature-controlled (68–79°F), humidity-controlled (30–70%), 12 hr light/dark cycle, with provided food and water *ad libitum*. Prior to surgery, animals were randomized into experimental groups and initially anesthetized with 3% isoflurane followed by a Ketamine (100 mg kg^−1^): Xylazine (10 mg kg^−1^) cocktail and Buprenorphine SR (1 mg/kg) administered subcutaneously. An injury of 1.5 mm diameter, spanning the full thickness of the mucosal tissue, was made between the 3rd and 4th rugae using a cauterizer (Bovie Medical Corportaion). 3.8 µL hydrogel scaffolds (blank, UT-EV, SMase-EV) were implanted at the site immediately after ONF injury (Figure S7) and secured using Tegaderm tissue adhesive. On day 7 the palate tissue was harvested; mice were euthanized via CO_2_ asphyxiation and cervical dislocation. The entire hard palate mucosa tissue was excised using a scalpel with #11 straight, angled blade (Integra Miltex 4-111) and immediately placed on ice in 3% FBS/PBS for further processing.

### Flow Cytometry Analysis of Murine Palate Tissue

Flow cytometry experiments were performed using samples from 3-4 male C57BL/6J mice per treatment group. Palatal mucosal tissue was freshly collected and enzymatically dissociated in a digestion buffer containing Collagenase II (5500 U/mL, Thermo Fisher Scientific, #17101015) and Dispase II (2.5 U/mL, Sigma-Aldrich, #D4693-1G) for 60–80 minutes at 37 °C. Following digestion, single-cell suspensions were rinsed with ice-cold PBS supplemented with 3% FBS, centrifuged for 5 minutes, and the resulting pellets were retained. Cells were blocked using Mouse Fc Block (BD Biosciences, #101301) and stained with an optimized antibody panel as previously described [41]. After staining, samples were fixed in 4% paraformaldehyde and stored overnight at 4 °C in FACS buffer prior to acquisition. Data collection was carried out on a five-laser Cytek Aurora cytometer, and downstream analyses, including gating strategies and Uniform Manifold Approximation and Projection (UMAP) visualization (n_neighbor = 15 and min_distance = 0.4), were conducted using the OMIQ.ai platform.

### Quantification and Statistical Analysis

Normality and variance were assessed by Shapiro-Wilk, F-test, or Spearman’s tests. Statistical significance was defined as p < 0.05 and evaluated using unpaired t-tests, one-way ANOVA with Tukey’s post-hoc tests for multiple comparisons, or two-way repeated measures ANOVA when appropriate. If assumptions were violated, nonparametric tests (Unpaired t-test with Welch’s correction, Mann-Whitney, or Kolmogorov-Smirnov) were used. All statistical analyses were performed with GraphPad Prism version 10.3.0 for Windows (GraphPad Software, Boston, Massachusetts USA, www.graphpad.com). All information pertaining to error bars, p-values, statistical tests, and sample sizes are reported in the figure legends.

## RESULTS

### Lipid Raft Organization and Sphingolipid Distribution Are Significantly Altered by SMase

We assessed whether SMase influences MSC membrane dynamics, particularly lipid rafts and associated sphingolipids (Figure 1A). Immunofluorescence staining of CD63-positive compartments (green) and ganglioside GM1 (red) showed that SMase-treated MSCs retained comparable CD63 expression levels (Figure 1B–C) yet exhibited a significant increase in GM1 area (Figure 1D). This expansion suggests SMase reshapes the PM microdomain landscape, a phenomenon that may have implications for vesicle budding and cargo sorting. To further investigate these shifts, we performed untargeted sphingolipid profiling. The corresponding z-score heatmap (Figure 1E) revealed broad variations in lactosylceramide and ganglioside species as these glycosphingolipids were found in lipid rafts between untreated (UT) and SMase-treated cells, with some species (e.g., LacCer, GM3) notably elevated under SMase treatment. In addition, conditioned media was collected from NBD-labeled sphingomyelin (NBD-SM) treated MSCs to understand secreted lipid dynamics. These SMase-treated samples were enriched with NBD-Cer and NBD-glucosylceramide (Figure S1 A-B) with the max concentration noted at 3 and 9 hours, respectively. While NBD-fatty acids concentration was unaltered and NBD-SM was decreased in SMase-treated samples (Figure S1C-D). These findings were supported by net enzymatic exchange fluxes models (Figure S1E-H). Untargeted lipidomics confirmed cellular lipid alterations showcasing a shift in Cer abundance from 7.99% to 30.48% between UT and SMase treatment groups (Figure 1H). Similarly, this stark trend was also observed between EV groups with 3.39% Cer abundance in UT and 48.91% in SMase-treated (Figure 1I). Next, we wanted to assess if SMase treatment would alter MSC biophysical properties, thus we conducted atomic force microscopy on UT and SMase-treated MSCs. We saw a significant increase in MSC stiffness following SMase treatment (Figure 1F-G), suggesting underlying structural changes throughout the cells are attributed to changes in lipid profile. Collectively, these data highlight that SMase reorganizes lipid raft domains and modifies sphingolipid composition while preserving core exosomal membrane protein, CD63, providing a mechanistic basis for downstream effects on vesicle biogenesis, cargo sorting and cell signaling.

**Figure 1.**
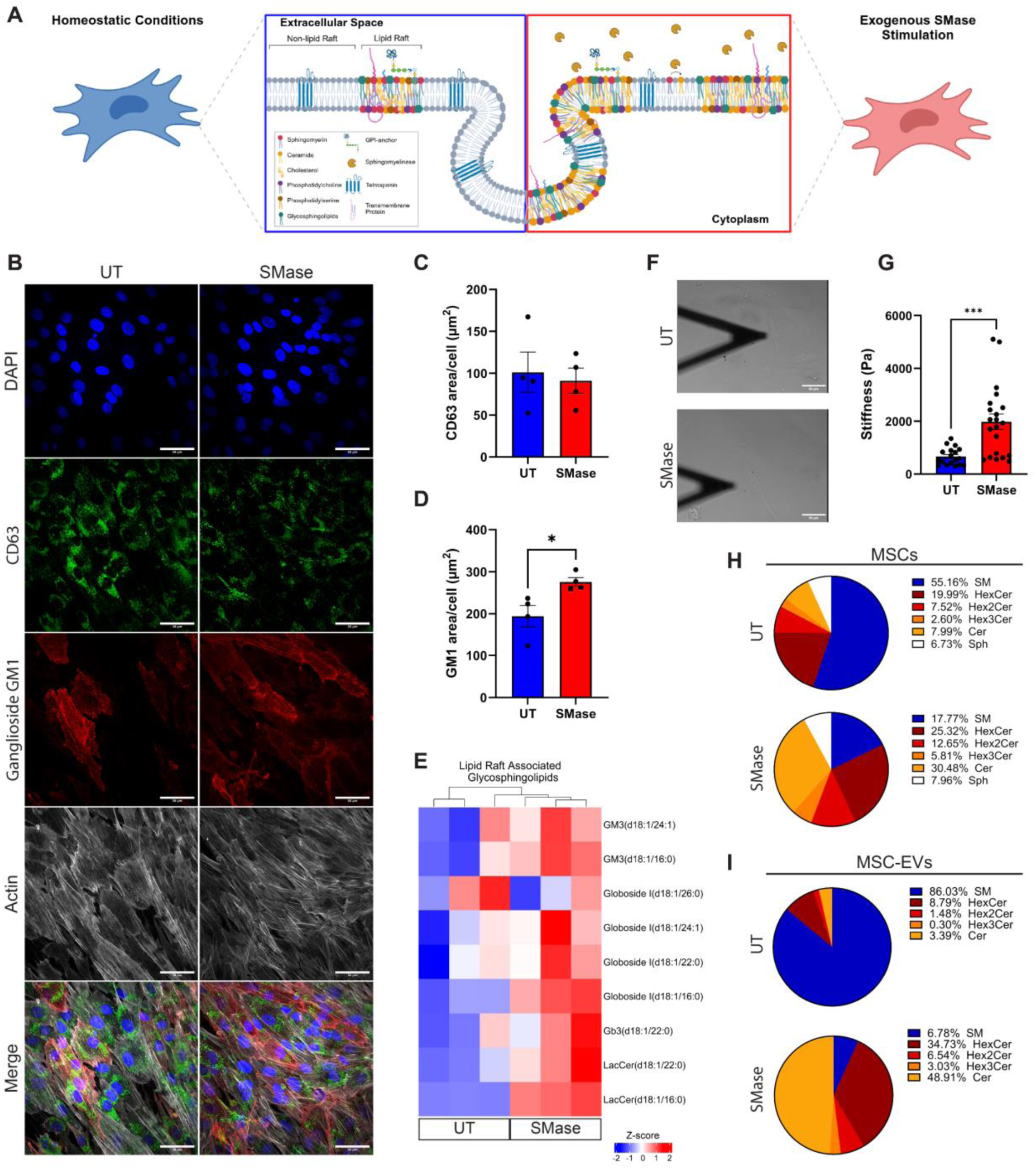
SMase treatment alters MSC lipid raft organization and sphingolipid distribution. (A) Graphical representation of MSC lipid microdomain alterations following 24-hour treatment of exogenous SMase. (B) Representative immunofluorescence images showing nuclear staining (DAPI, blue), CD63-positive compartments (green), Ganglioside GM1 (red), and F-actin cytoskeleton (gray) in untreated (UT) and SMase-treated MSCs (scale bars = 50 μm); n=4. (C) Quantification of CD63-positive area per cell between treatment groups. (D) Analysis of lipid raft area per cell in SMase-treated MSCs. (E) Z-score heatmap showing differential expression of glycosphingolipid species between UT and SMase-treated MSCs, with red indicating increased and blue indicating decreased; n=3. (F) Representative images and (G) quantification from atomic force microscopy (AFM) analysis of MSCs treated with or without SMase; n=19-21. (H) Annotated sphingolipid composition of MSCs and (I) EVs in UT and SMase-treated groups; n=3. Data presented as mean ± SEM. Statistical significance determined by unpaired t-test or Mann-Whitney test; *p < 0.05, ***p <0.001.

### SMase Enhances MSC-Derived EV Yield and Alters their Surface Marker Profile

To examine whether SMase influences extracellular vesicle (EV) characteristics and production, we performed nanoparticle tracking analysis (NTA), transmission electron microscopy (TEM), zeta potential, and biochemical assays on EVs harvested from UT or SMase-treated MSCs (Figure 2A). TEM micrographs verified the presence of typical EVs with cup-shaped morphology in both groups, although SMase-derived EVs appeared slightly more heterogeneous in size (Figure 2B). Representative NTA size distribution profiles (Figure 2C) and average diameter (Figure 2E) indicated a significant shift in the EV population size with increased mean diameter following SMase treatment. When assessing particle concentration, we detected a significant log-fold increase in the SMase stimulated group (Figure 2D). Furthermore, the zeta potential (surface charge) was significantly altered in the SMase-treated condition (Figure 2F), indicating possible rearrangements in membrane lipid composition. Quantification of EV protein revealed similar total protein concentration among EVs isolated from treated and non-treated MSCs (Figure 2G). RNA was quantified, uncovering the SMase EV group had a trending increase in total RNA (Figure 2H), suggesting SMase stimulation promotes alterations in the molecular packaging of EVs. Lastly, flow cytometric analysis confirmed that both UT and SMase-derived EVs expressed classical EV markers (CD9, CD63, CD81) (Figure S2A-C), although their relative abundance and per-particle marker levels varied between groups (Figure 2I–L). Collectively, these data demonstrate that SMase treatment increases EV yield, modifies EV surface charge, and alters marker composition, findings that point to membrane remodeling as a potential mechanism underlying these changes.

**Figure 2.**
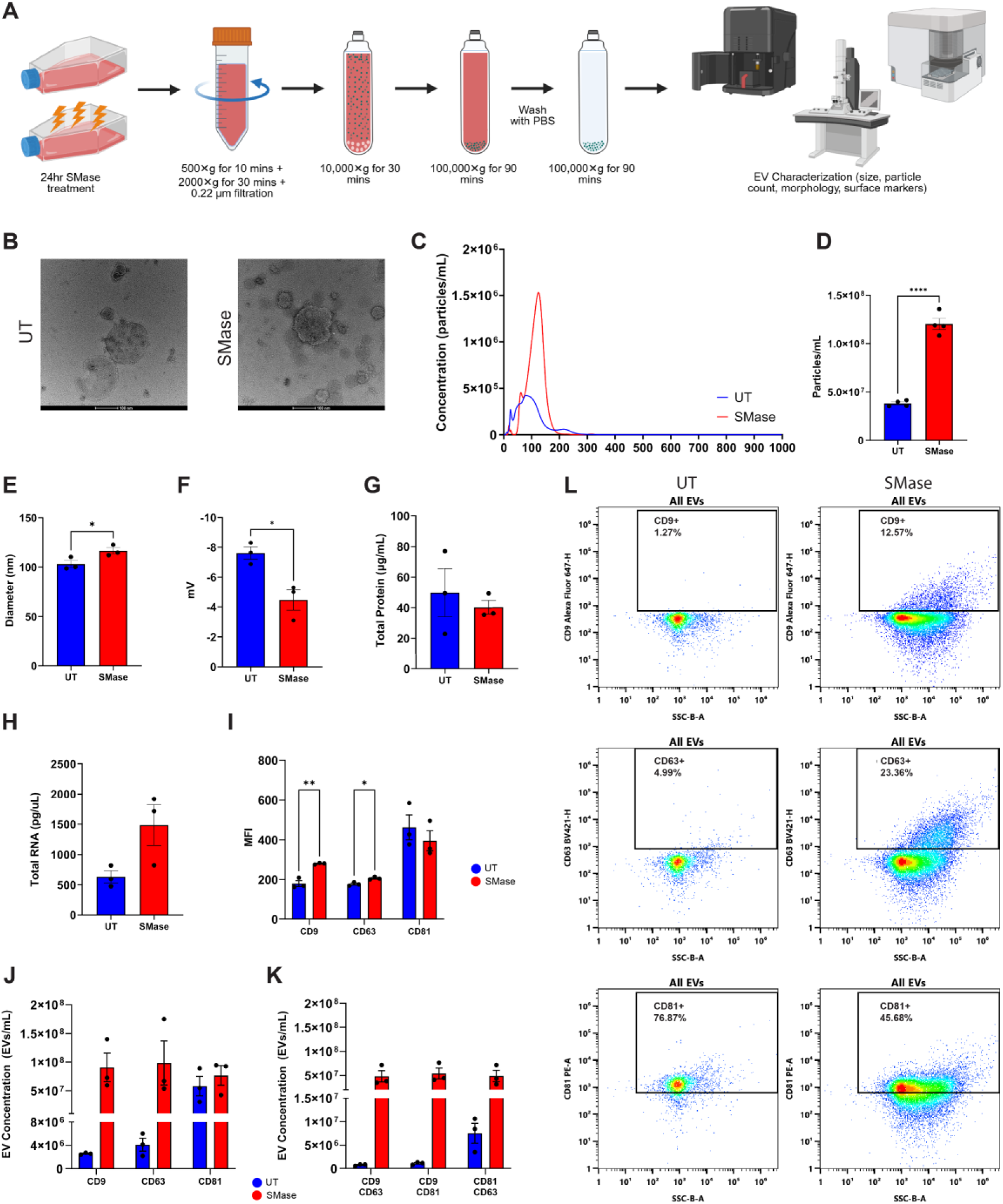
SMase treatment enhances MSC-derived EV yield and alters their surface marker profile. (A) Experimental workflow of EV isolation and characterization. (B) Transmission electron microscopy (TEM) images showing morphology of isolated EVs from both groups (scale bar = 100 nm). (C) Representative nanoparticle tracking analysis (NTA) size distribution profiles of EVs isolated from untreated (UT) and SMase-treated MSCs. (D) Quantification of particle concentration and (E) average particle diameter between EVs derived from UT and SMase treated MSCs. (F) Zeta potential measurements between UT and SMase-derived EVs. (G) Quantification of EV protein concentration. (H) Total RNA concentration from treated and untreated MSC-derived EVs. (I) Quantification of classical EV markers (CD9, CD63, CD81) showing differential MFI between groups. (J) Analysis of vesicle concentration in single positive EV populations and (K) double positive populations. (L) Representative flow cytometry plots showing surface marker expression on EVs. Data presented as mean ± SEM. Statistical significance determined by unpaired t-tests or one-way ANOVA; *p < 0.05, **p < 0.01, ***p < 0.0001; n=3.

### SMase Induces Distinct Lipid Signatures in MSCs and Their Derived EVs

To gain deeper insight into how SMase shapes lipid networks, we performed comprehensive lipidomic profiling of both MSCs and their EVs (Figure 3A). Hierarchical clustering and volcano plots (Figure S3A–B and Figure S4A) revealed pronounced differences in lipid expression between UT and SMase-treated MSCs, with 214 features significantly up-regulated and 156 down-regulated. Lipid pathway enrichment analysis (LiPEA) highlighted “glycerophospholipid metabolism”, “sphingolipid signaling pathway”, and “glycosylphosphatidylinositol (GPI)-anchor biosynthesis” pathways as notably impacted (Figure S3C). Following SMase stimulation, MSC membranes showed ceramide enrichment and complementary shifts in sphingolipids. A parallel analysis of their secreted EVs (Figure S3D–E and Figure S4B) revealed 91 up-regulated and 243 down-regulated features, underscoring EV lipid composition recapitulating the cellular remodeling present at the time of release. LiPEA results (Figure S3F) showed pathways involving “sphingolipid signaling pathway”, “sphingolipid metabolism”, and “retrograde endocannabinoid signaling” to be altered following SMase treatment.

**Figure 3.**
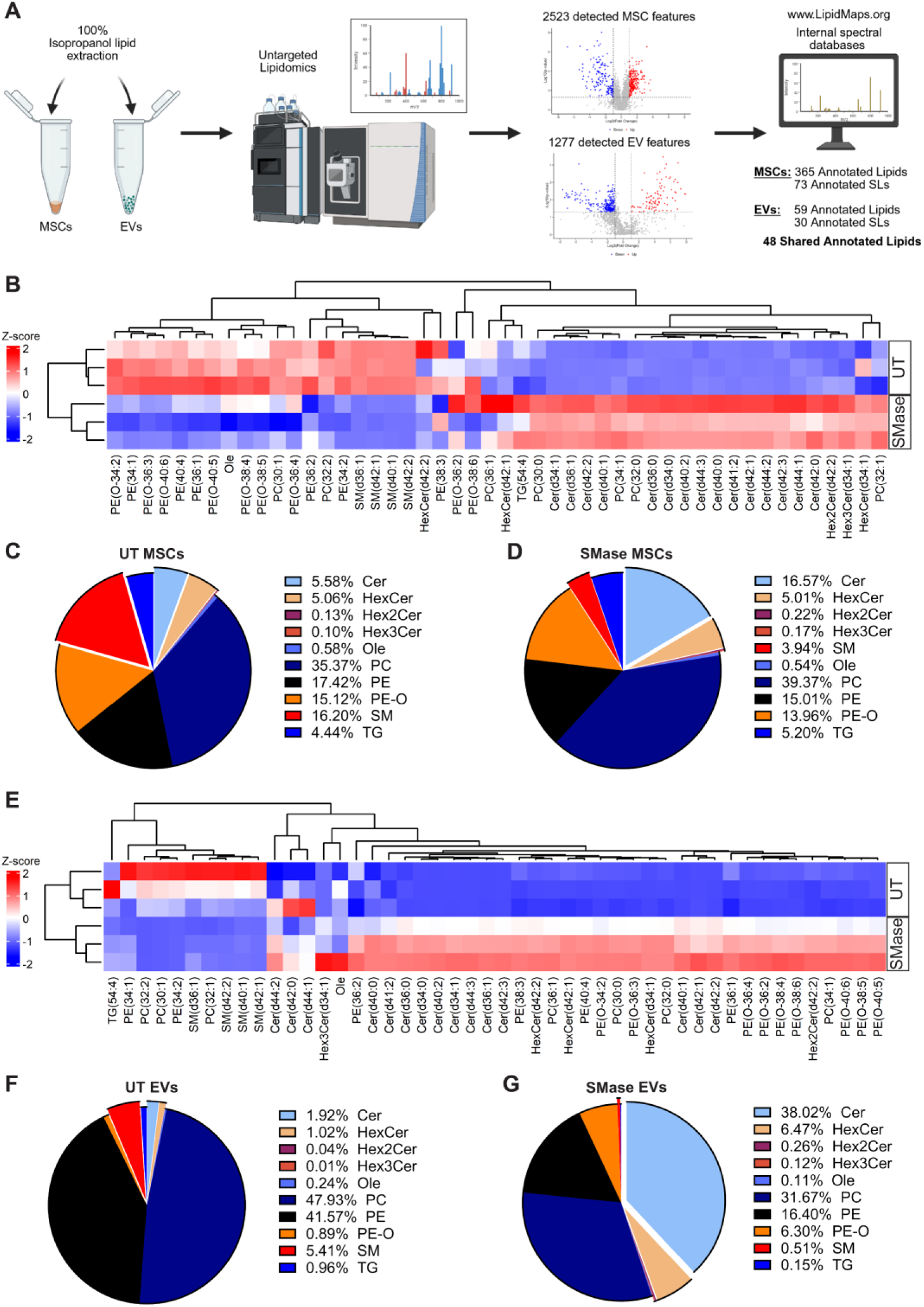
SMase treatment induces distinct lipid class remodeling in MSCs and their derived EVs with enhanced ceramide enrichment. (A) Graphical depiction of lipid isolation, quantification, and data analysis workflows. (B) Hierarchical clustering heatmap showing differential lipid expression of shared annotated lipids in untreated (UT) and SMase-treated MSCs (blue: downregulated; red: upregulated). Pie charts showing relative distribution of major lipid classes: (C) UT MSC (D) SMase-treated MSC. (E) Hierarchical clustering heatmap demonstrating shifts in lipid profiles of shared annotated lipids between UT and SMase-derived EVs (blue: downregulated; red: upregulated). Pie charts showing relative distribution of major lipid classes in (F) UT EVs and (G) SMase-treated EVs. n=3.

We explored more deeply into specific lipid classes and plotted abundances of individual annotated ceramides, sphingomyelins, glycerophospholipids, and other major categories (Figure S4C–F). Compared with UT controls, SMase-treated MSCs and their derived EVs showed trending log increases in several ceramide acyl chains such as, Cer (34:0), (34:1), (36:0), (36:1) (Figure S4C, E), consistent with the enzymatic conversion of SM to Cer. In parallel, phosphatidylcholine (PC) and phosphatidylethanolamine (PE) distributions in MSCs were similar between groups (Figure S4D), but trending log-fold shifts were noticed in EVs (Figure S4F). Z-scored heatmaps of the 48 shared SLs and phospholipids between both MSCs (Figure 3B) and their derived EVs (Figure 3E) identified shared remodeling of lipids. Pie chart comparisons provided an overview of how these alterations reshaped the lipidome: SMase-treated cells (Figure 3C-D) and their EVs (Figure 3F-G) exhibited a marked enrichment in Cer species and EV phospholipid levels (PE and PC) decreased relative to UT EV samples. These findings highlight SMase treatment facilitates lipid class remodeling in both the cellular and EV compartments, potentially influencing membrane curvature, vesicle budding, and vesicle cargo sorting.

### Proteomic Remodeling Highlights Enrichment of RNA Processing, RNA-Binding, and Membrane Organization Pathways

Beyond lipid changes, we investigated whether SMase treatment also affects MSC protein expression profiles. Hierarchical clustering of differentially expressed proteins (Figure S5A) demonstrated clear segregation between UT and SMase-treated MSCs, with 34 downregulated and 143 upregulated features (Figure S5B). Focused analysis of selected clusters (Figure 4B) revealed upregulation of proteins involved in vesicular trafficking (Rab6a, Rab11b, and Rab14), RNA-binding proteins (HNRNPDL, HNRNPA2B1, and YBX1), GPI-anchor synthesis (PIGG), and immunomodulation (APOE and HMGB1). Gene ontology (GO) enrichment confirmed that SMase treatment enhances MSC molecular function pathways linked to “RNA binding” and “structural molecule activity” (Figure S5E). Biological processes (Figure S5C) observed significant enrichment in “translation”, “RNA processing”, and “ribosome biogenesis”. The cellular component category (Figure S5D) further corroborated increased pathways associated with “vesicle” and “ribonucleoprotein complex”. Taken together, these proteomic data imply that SMase not only affects sphingolipid metabolism but also orchestrates changes in the molecular machinery underlying RNA processing and membrane organization. This coordinated reprogramming of lipid and protein networks likely converges to facilitate EV production and cargo packaging, in alignment with the heightened EV release observed (Figure 2).

**Figure 4.**
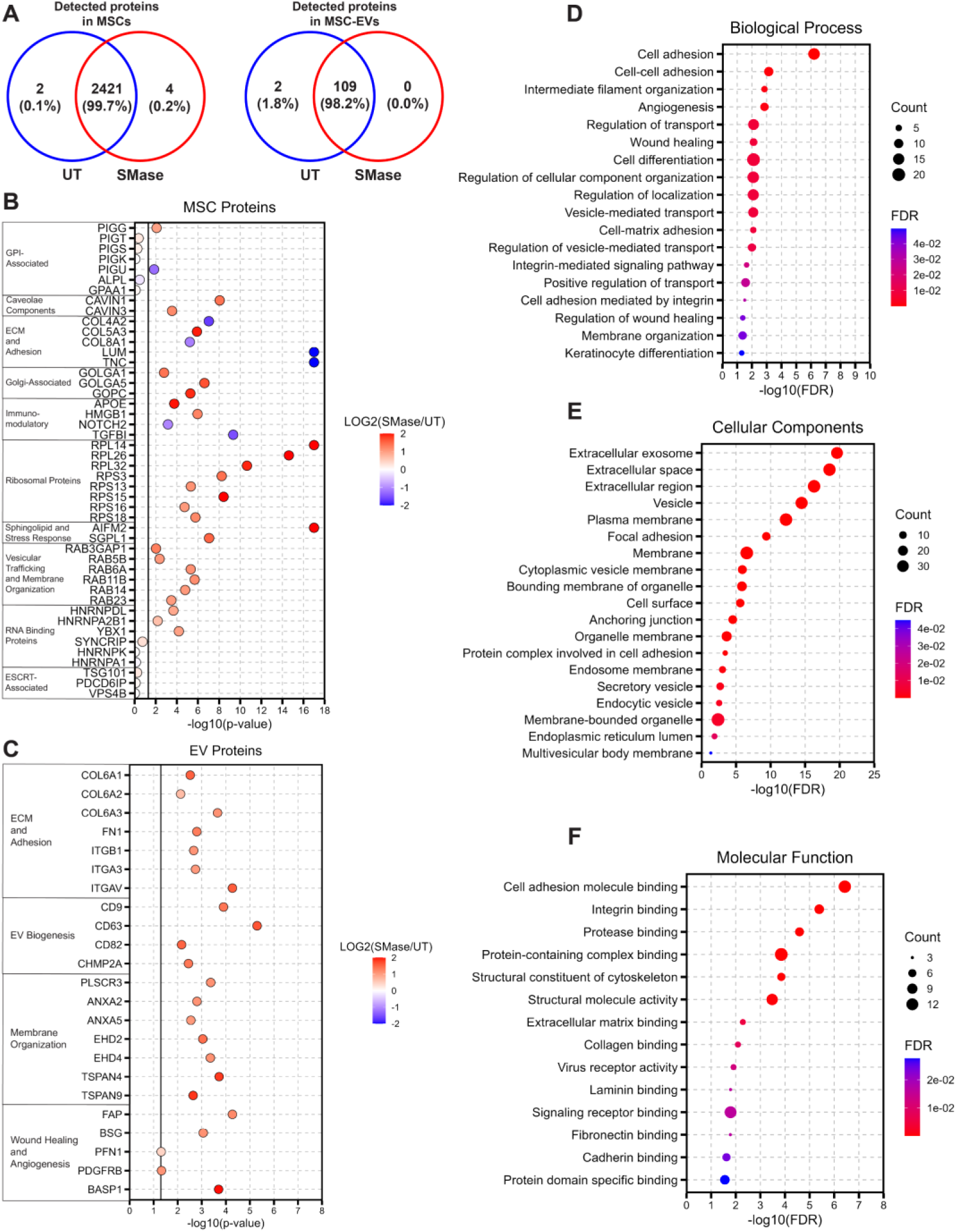
SMase treatment alters protein cargo composition with enrichment of RNA-binding, angiogenic, and wound healing-associated proteins. (A) Venn diagrams outlining the detected features in MSCs and MSC-EVs. (B-C) Dot plots of key protein clusters in (B) MSCs and (C) MSC-EVs; data to the right of the black line indicates p < 0.05, color intensity represents log_2_(fold change) value. (D-F) EV protein gene ontology analysis demonstrating significantly enriched terms in (D) biological processes, (E) cellular components, and (F) molecular functions. GO analysis statistical significance determined by false discovery rate-adjusted p-values < 0.05; dot size indicates protein count; color intensity represents FDR value. n=3.

### SMase Induces Proteomic Remodeling in EV Cargo and Vesicular Trafficking Proteins

To investigate how SMase modifies the proteomic landscape of MSC-derived EVs, we performed shotgun proteomics on isolated EVs from UT or SMase-treated MSCs (Figure 4). Hierarchical clustering revealed a clear separation between UT and SMase EV proteomes (Figure S6A), with 35 proteins upregulated and 6 proteins downregulated upon SMase treatment (Figure S6B-C). A focused analysis of functional clusters showed significant enrichment in wound healing and angiogenic-associated proteins (FAP and BSG), vesicular biogenesis components (CD9, CD63, CD82, and CHMP2A), and membrane organizers (TSPAN4, TSPAN9, and PLSCR3) (Figure 4C). Corresponding GO terms underscored significant increases in pathways associated with “angiogenesis”, “vesicle-mediated transport”, “wound healing”, “protein-containing complex binding”, and “membrane organization” (Figure 4D-F). Together, these data suggest that SMase prompts a targeted recruitment of proteins into EVs that could bolster EV cargo packaging through altered vesicular protein trafficking and membrane-associated processes.

### SMase Reshapes MSC and EV miRNA Profiles, Enhancing Immunomodulatory Pathways

We next explored whether SMase-driven changes in EV protein cargo are paralleled by shifts in miRNA composition (Figures 5A). Differential expression analyses identified 8 upregulated miRNAs in SMase treated MSCs, including miR-143-3p, miR-27a-5p, and miR-21-3p, while similarly 8 miRNAs (e.g., miR-335-5p, miR-29b-3p) were downregulated (Figure 5B). Targets of these differentially expressed miRNAs (DEmiRNAs) revealed pathways linked to “endocytosis”, “MAPK signaling pathway”, and “protein processing in endoplasmic reticulum” (Figure 5C). Differential expression analysis in EVs resulted in 20 upregulated (e.g., miR-23b-3p, miR-31-5p, and miR-191-5p) and 13 downregulated (e.g., miR-34a-5p, miR-138-5p) miRNAs within the SMase EVs (Figure 5D). In alignment with the MSCs, targets of these DEmiRNAs are associated to pathways involving “MAPK signaling pathway” and “endocytosis”, as well as “TNF signaling pathway” and “efferocytosis” (Figure 5E). In addition, EVs derived from SMase-treated MSCs displayed 203 specific miRNAs compared to 24 unique miRNAs in UT MSC-derived EVs (Figure 5F). When comparing the miRNA profiles to that of the parental cells, UT MSC-derived EVs had 12 uniquely detected miRNAs accounting for 1.4%, while 41 miRNAs were exclusive to SMase EVs when compared to SMase-treated MSCs (Figure 5G).

**Figure 5.**
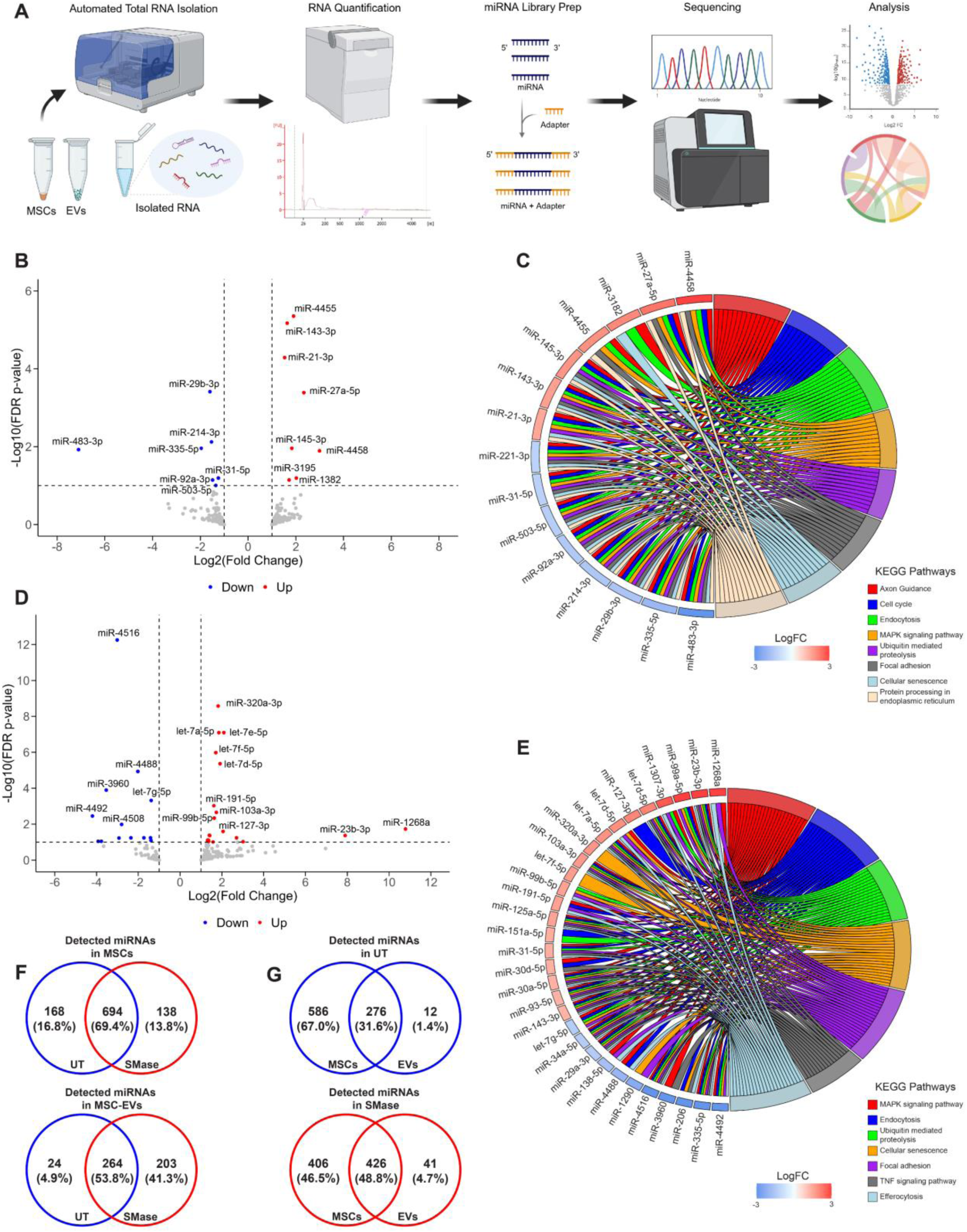
SMase treatment enhances selective miRNA sorting into MSC-derived EVs. (A) Workflow of miRNA isolation, quantification, and data analysis. (B)Volcano plot demonstrating global miRNA expression changes in MSCs. (C) Chord plot depicting the relationship between MSC DEmiRNAs (blue: downregulated; red: upregulated) and their target KEGG pathways. (D) Volcano plot indicating significantly altered miRNAs in MSC-derived EVs following SMase treatment. (E) Chord plot indicating associations among MSC-EV DEmiRNAs and their target KEGG pathways, fold change values indicated by blue (downregulated) and red (upregulated). (F-G) Venn diagram outlining detected miRNA distribution among (F) UT vs. SMase in each sample type and (G) parental cells vs. derived EVs for each treatment group. Statistical significance determined by false discovery rate-adjusted p-values < 0.1 and |FC| > 2. n=3

### SMase-Derived EVs Enhance Angiogenic Potential and Immunomodulatory Effects

We tested whether these shifts in EV composition translate into functional benefits (Figure 6). Using an *in vitro* angiogenesis assay, we observed media supplemented with SMase-derived EVs trended to increased tubular network formation compared to UT EVs or control media, evidenced by a trend of greater numbers of nodes, meshes, and branches (Figure 6A–E). In parallel, we assessed the immunomodulatory capacity of SMase-treated MSC-derived EVs in an *in vitro* macrophage activation model. When LPS-stimulated macrophages were co-incubated with SMase-derived EVs, TNF-α production was significantly reduced relative to LPS-treated (control) macrophages. Macrophages treated with SMase-derived EVs also showed a trend of TNF-α reduction compared to UT EVs (Figure 6F–G). In addition, a pilot *in vivo* study was conducted using a critical murine oral wound defect model. Palatal tissue was explanted 7 days following injuring and/or treatment followed by cellular phenotyping conducted via single cell flow cytometry (Figure S7). Single cell flow cytometry data across all groups were pre-gated and all events from CD45^+^CD11b^-^lymphoid cells were used to construct a uniform manifold approximation and projection (UMAP) (Figure 6H). The UMAP surface maker expression is represented from low (black) to high (copper), with all groups overlayed. Cell density is displayed ranging from low (blue) to high (red) for injured, hydrogel, UT-EV hydrogel, and SMase-EV hydrogel groups. UMAP visualized cell density of CD8+, CD19+, and CD11c+ events are shown to be lower in both UT-EV and SMase-EV hydrogel groups compared to injury and hydrogel control groups. Lymphoid cell populations were identified (Figure 6I) showing mean changes in T-cell, B-cell, and Dendritic cell distribution based on treatment group. Reduced cellular distribution of CD3^+^ and CD8^+^ T-cells was found subsequently in the SMase-derived EV hydrogel treated mice compared to UT-EV hydrogels, while no change was found in CD4^+^ T-cell distribution (Figure 6H-K). Together, these results suggest that SMase-induced changes in EV cargo (both proteomic and miRNA) may confer enhanced pro-angiogenic and anti-inflammatory effects.

**Figure 6.**
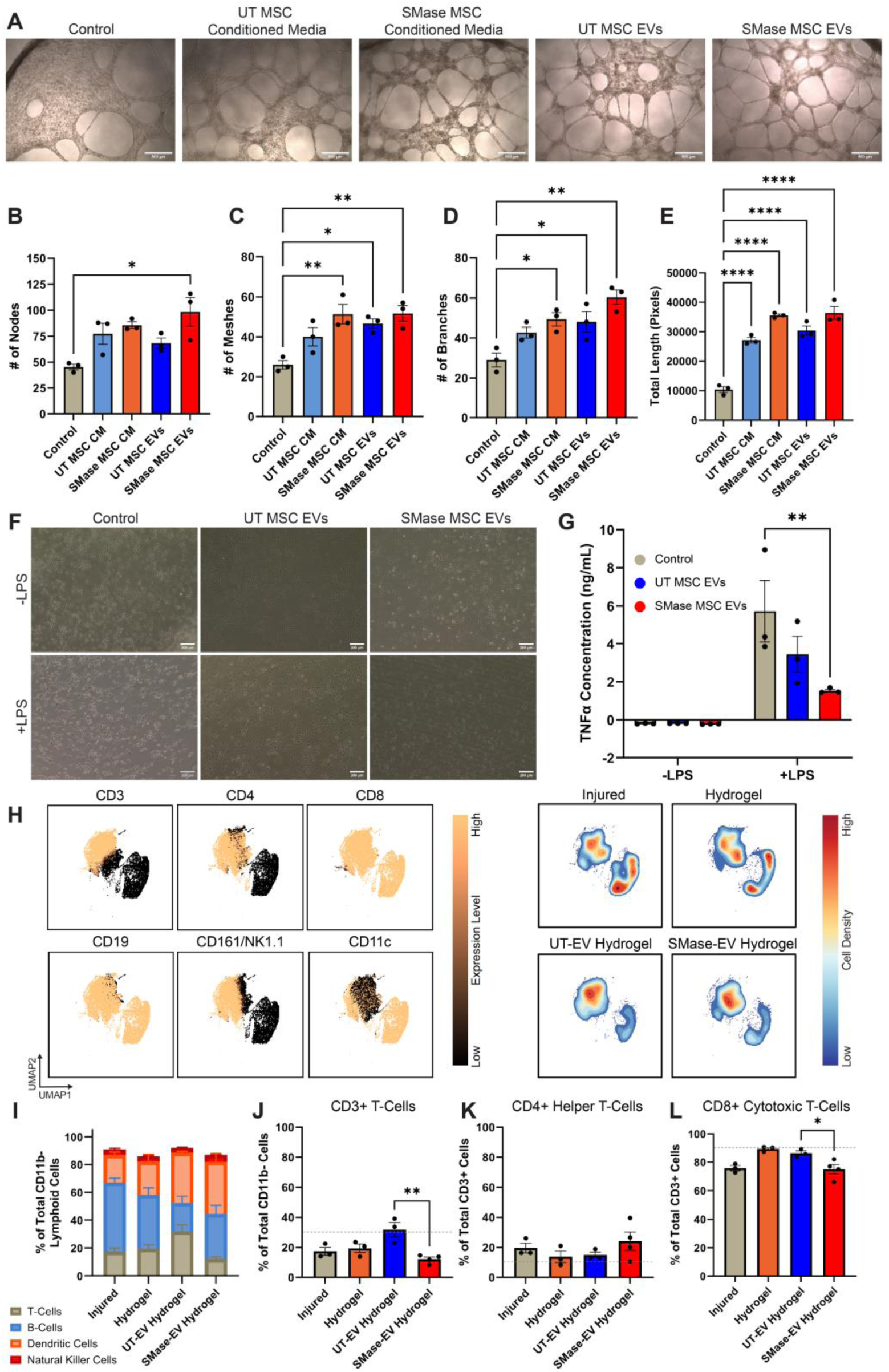
SMase-treated MSC-derived EVs enhance angiogenic potential and immunomodulatory function. (A) Representative phase-contrast images showing microvascular network formation in response to control media, untreated (UT) and SMase-treated MSC conditioned media, and their respective EVs (scale bars = 500 μm). (B-D) Quantitative analysis of network formation parameters showing enhanced (B) number of nodes, (C) number of meshes, (D) number of branches, and (E) total length in all conditions. (F) Representative images of macrophage response to LPS stimulation in the presence of UT or SMase MSC-derived EVs (scale bars = 200 μm). (G) Quantification of TNF-α production following treatment and LPS-stimulation in macrophages. Single-cell flow cytometry performed on wounded tissues collected from mice treated with hydrogels, UT-EV hydrogels, or SMase-EV hydrogels at day 7 post-injury. (H) UMAP projection of single-cell flow cytometry data illustrating lymphoid cell marker expression intensity from low (black) to high (copper) and corresponding cell density from low (blue) to high (red). (I) Stacked bar graph depicting quantified CD11b^-^ immune cell populations. T-cell populations were identified and classified as (J) CD3⁺ T-cells, (K) CD4⁺ helper T-cells, (L) CD8^+^ cytotoxic T-cells. Data presented as mean ± SEM. Statistical significance indicated as *p < 0.05, **p < 0.01, ***p < 0.001, ****p < 0.0001; n=3-4.

## DISCUSSION

MSC-EVs have emerged as a promising cell-free therapeutic platform due to their inherent immunomodulatory and pro-regenerative properties. However, challenges in scaling production and heterogeneity have limited their clinical translation. Here, we explored a lipid-centric approach to engineering EVs by leveraging the enzymatic activity of exogenous SMase, a key mediator of Cer biosynthesis and membrane curvature dynamics. Our findings demonstrate SMase treatment of MSCs offers a compelling strategy to enhance EV production while reshaping vesicle content by modifying cargo sorting of potent proteins and miRNAs. Importantly, our previous data confirm that controlled SMase stimulation of MSCs is well-tolerated, showing no evidence of cytotoxicity, altered proliferation, or loss of viability, consistent with previous reports indicating that moderate modulation of sphingolipid metabolism preserves cellular integrity while enhancing vesicle secretion [32, 34, 42]. This work builds on and extends the current understanding of ceramide-driven EV biogenesis and provides insights into how targeted lipid remodeling can orchestrate downstream effects on vesicle formation, molecular composition, and paracrine function.

The expansion of lipid raft domains and sphingolipid redistribution observed in this study underscore a potential mechanism through which SMase promotes cargo trafficking and ceramide-dependent EV biogenesis. Prior reports highlighted ceramide’s role in membrane curvature and intraluminal vesicle formation [15, 22, 23, 25, 26]. Notably, it has been reported that RNAi knockdown of SMPD2/3 led to significant reduction in EV production [23]. Studies have also suggested lipid raft microdomains may be integral in the formation of therapeutic EVs [17, 19]. Our findings bolster these precedents, demonstrating exogenous SMase can accelerate Cer generation in both MSC and EV membranes, and in turn, promote large scale EV release through ceramide-dependent exosome formation [14, 43]. This is consistent with our earlier findings where SMase-licensed MSCs exhibited increased Cer and GlcCer and leading to redistribution of SM in cellular samples [34]. In addition to Cer synthesis, downstream lipid metabolism was identified with Cer preferentially converted into glycosphingolipids (HexCer, Hex2Cer, Hex3Cer, LacCer, and GM3) shown by the high levels of overall glycosphingolipids compared to sphingosine following SMase treatment. Notably, glycosphingolipids delivered to macrophages from EVs have been found to enhance their phagocytic function and induce suppression of T cells [44]. Additionally, SLs such as glycosphingolipids have been identified as essential metabolites involved in immune cell evasion of solid tumors [45] and the promotion of angiogenesis [46, 47], highlighting their potent role in immunomodulation and vascular remodeling.

To further probe putative cellular and exosomal modifications, we coupled lipidomics, proteomic and miRNA transcriptomic profiling and found SMase-mediated alterations in the MSC lipidome synergize with proteomic shifts that promote EV assembly and cargo loading. Notably, proteomic changes in parent cells reveal a concurrent upregulation of caveolae, vesicular trafficking, and RNA binding proteins, suggesting the remodeling of sphingolipid metabolism is accompanied by an intrinsic restructuring of the molecular machinery supporting enhanced lipid and protein processing. Specifically, Cavin1 is well known for its role in selectively sorting PM lipids predicated on head group and acyl chain to form anomalous nanostructures such as lipid rafts [48]. These structures have been reported as the initial budding and endosomal formation necessary for ESCRT-independent EV biogenesis. In MSCs, EVs formed from lipid raft microdomains require the presence of Cer confirmed by reduced GM1 and CD81 rich EVs following SMase inhibition with GW4869 [17]. Within our study, we notice unchanged ESCRT-associated proteins which facilitate vesicular trafficking and formation, while seeing increased activity of several small GTPase proteins known to also play a role in vesicular trafficking and release. Specifically, RAB23’s involvement in the endocytic pathway has been investigated and shown to facilitate PM endocytosis [49]. In addition, RAB11 has been implicated as a key regulator involved in the release of exosomes [50]. Tangential to vesicular trafficking proteins, several RNA-binding proteins (RBPs) involved in cargo sorting were differently expressed following SMase treatment. Following covalent bonding of small ubiquitin-like modifier proteins, HNRNPA2B1 has been recognized to bind and directly traffic miRNAs into exosomes [51]. In parallel, YBX1 has been shown to preferentially bind and sort miRNAs into HEK293T-derived exosomes [52]. This data reveals intriguing information worth further investigation to mechanistically understand the link between sphingolipid metabolism and RNA-binding protein recruitment.

The selective enrichment of regulatory proteins in SMase-treated MSCs is mirrored with the observed changes in SMase-induced EV miRNA composition. These coordinated shifts likely improve selective miRNA encapsulation of immunomodulatory miRNAs and bolster the molecular complexity of the resultant EV cargo. Similar to recent findings in other cell types [31], the enhanced packaging of specific therapeutic miRNAs suggests that membrane reorganization influences cargo sorting mechanisms [20, 32, 53]. Short non-coding RNAs such as miR-23b-3p and miR-191-5p are known for their ability to regulate immune cells. In particular, when miR-23b-3p was loaded into MSC-derived exosomes targeting for alveolar macrophages and delivered into mice with acute lung injury, there was a reduction in M1-like macrophages mediated though Lpar1-NF-kB pathway regulation [54]. Similarly, miR-191-5p has been shown to activate an M2-like macrophage phenotype through inhibition of SOCS3 expression [55]. The notion of co-regulated membrane remodeling processes and cargo packaging is further supported by altered surface properties (e.g., zeta potential, membrane bound proteins, and lipids). Similar findings in other model systems indicate that membrane composition influences EV budding, cargo loading, and eventual uptake by recipient cells [9, 15].

Functionally, the enhanced immunomodulatory and pro-angiogenic activity of SMase-derived EVs suggests that Cer-fueled lipid raft reorganization may yield more potent EVs. We detected elevated levels of pro-angiogenic factors and immunomodulatory miRNAs (FAP and BSG, and miR-31-5p), paralleling previous work that highlights Cer-driven modifications to exosome cargo [22, 31] and MSC-EV therapeutic applications [12, 17, 19]. In addition to the identification of these factors, functional *in vitro* testing of SMase-induced EVs treating activated macrophages showed a trend in reducing TNF-α production, as recent clinical studies have highlighted the importance of this pathway in inflammatory conditions [13, 56]. Moreover, the trending increased formation of HUVEC tubular networks aligns with emerging evidence that EV membrane composition influences their uptake and functional impact on recipient cells [3, 15, 57]. Therapeutic function was also seen *in vivo* when delivering SMase-EVs into local critical oral wound beds within mice. The SMase-EVs significantly altered CD3^+^ and CD8^+^ T-cell distribution following 7 days post injury and implantation. The trends observed by mean increased tube formation in endothelial cell cultures mean decreased pro-inflammatory cytokine output in macrophages stimulated with SMase-EVs, and reduced cytotoxic T-cell populations indicate that these enzymatically modified vesicles could offer meaningful therapeutic advantages in tissue repair and immune regulation. These findings support broader trends in regenerative medicine, where EV-based therapies have shown promise in inflammatory disease, cardiovascular repair, and wound healing [12, 53, 58]. Our observation of enhanced angiogenic responses is especially notable, as robust vascularization is key to functional tissue recovery in ischemic and chronic injury contexts [10, 57]. The parallel immunosuppressive properties could expand the application of SMase-modified EVs to autoimmune disorders or graft-versus-host scenarios where dampening pathogenic inflammation is a primary therapeutic goal [3, 11].

This work exemplifies an alternate approach to EV engineering by directly harnessing exogenous SMase to modify lipid raft composition. Unlike genetic modifications, which may raise safety or regulatory concerns, the use of exogenous enzymes presents a relatively straightforward and potentially more transient method of augmenting EV yields. Moreover, our multi-omic characterization provides a rich mechanistic overview of how SMase reconfigures MSC architecture at multiple levels: lipid, protein, and miRNA, offering valuable data for the rational design of future EV-based therapeutics [59, 60]. Taken together, our findings highlight the potential of exogenous SMase as a tool to bolster the intrinsic paracrine potency of MSC-derived EVs. By specifically targeting sphingolipid pathways, we showed membrane remodeling can enhance EV biogenesis, tune cargo composition, and potentially enhanced angiogenic and immunomodulatory functions. The preservation of key EV markers while enhancing therapeutic cargo and EV production suggests SMase treatment could provide a scalable method for producing more potent EVs [16]. Such a strategy aligns with emerging trends in cell-free therapeutics, where modular control of vesicle composition aims to optimize tissue targeting, cargo stability, and functional efficacy [12, 13]. Beyond MSCs, this enzymatic approach could potentially be applied to other cell types, expanding the repertoire of EV-based therapies across various fields [4, 16]. Ultimately, leveraging sphingolipid metabolism to fine-tune EV properties represents an exciting frontier for precision-engineered cellular secretomes with broad therapeutic potential.

### Limitations of the study

While our *in vitro* assays demonstrate promising angiogenic and immunomodulatory activity of SMase-derived EVs, the sample size is low and increased sample populations could bolster the trends seen. In addition, these results may not fully predict *in vivo* or clinical responses. Additional animal models will be important for evaluating biodistribution, stability, and safety. The long-term stability of these engineered EVs, along with potential off-target effects of exogenous SMase on non-EV related pathways, remains uncharted. Our work was conducted with a single MSC donor and tissue source. Future studies should systematically examine donor- and tissue-specific variability. Although our multi-omic analyses provide insights into lipid remodeling and cargo enrichment, the precise molecular mechanisms by which SMase orchestrates cargo sorting remain to be fully elucidated. Addressing these open questions will be critical for translational development and scalable manufacturing of engineered EVs.

## ACKNOWLEDGEMENTS

This work was supported by the National Science Foundation Engineering Research Center on Cell Manufacturing Technologies (EEC1648035). D.C.S. and S.A.D. were supported by the NIH NIGMS-sponsored Cell and Tissue Engineering Biotechnology Training Grant (T32 GM145735 and T32 GM008433, respectively). The authors would like to thank the core facilities and personnel at the Parker H. Petit Institute for Bioengineering and Biosciences at Georgia Institute of Technology for shared equipment, services, and expertise: Samuel Moore, Dr. Rakesh Singh, Ludyanna Lebon, Shweta Biliya, Niama Djeddar, Erich Williams, and Dr. Danielle Scheff. The authors would also like to thank Chris Fleming with Cytek Biosciences for his assistance in developing EV flow cytometry methods. Figures were created using BioRender.com and Adobe Illustrator.

## AUTHOR CONTRIBUTIONS

D.C.S. and S.A.D. designed and conducted the experiments, analyzed the data, generated the figures, wrote, and revised the manuscript. H.Z. performed experiments, analyzed data, wrote, and revised the manuscript. A.Y.L. and T.A.A. performed experiments and data analysis. K.A.P assisted with figure creation. N.F.C performed data analysis, wrote, and revised the manuscript. A.I.T developed and oversaw experimental methods. E.A.B. supervised the project, generated figures, wrote, and revised the manuscript. Revising and approving the final version of manuscript: All authors.

