## Supplemental Materials for "Exogenous Sphingomyelinase Mediates MSC-derived EV Biogenesis and Enhances Potency via Repackaging of Molecular Cargo"

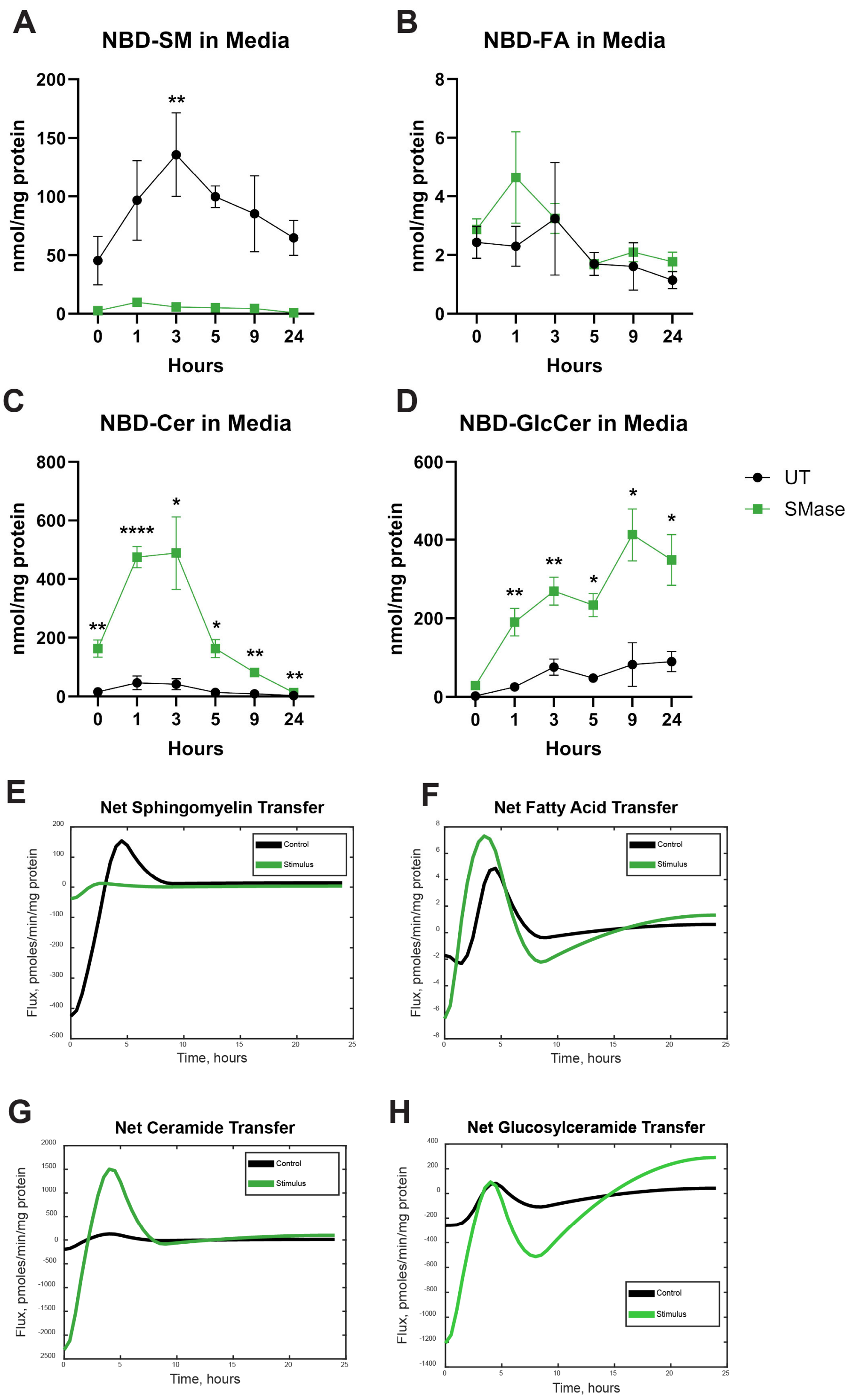


**Figure S1. NBD-lipid tracer identifies increased secreted ceramide and glucosylceramide in cell culture media following SMase stimulation, related to Figure 1.** Quantification of (A) NBD-fatty acid, (B) NBD-sphingomyelin, (C) NBD-ceramide, and (D) NBD-glucosylceramide concentration over time within cell culture media following NBD-SM addition in UT and SMase treated MSCs. Net enzymatic exchange fluxes in response to 24-hour SMase simulation for (E) Sphingomyelin, (F) Fatty acid, (G) Ceramide, and (H) Glucosylceramide. Data presented as mean ± SEM. Statistical significance determined by unpaired t-test with Welch's correction where appropriate; $^{*}p < 0.05$, $^{**}p < 0.01$, $^{***}p < 0.001$, $^{****}p < 0.0001$; n=3.

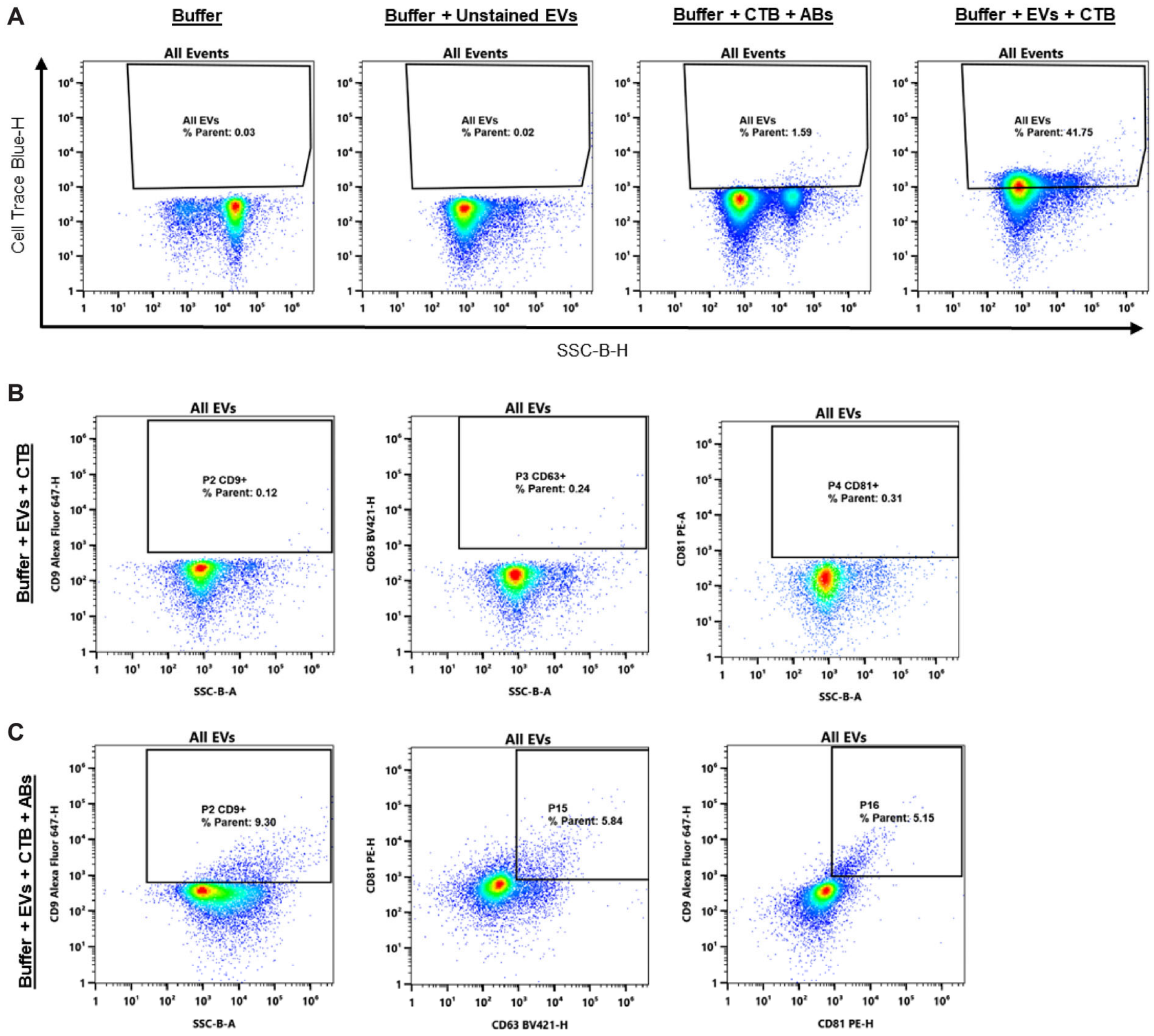


**Figure S2. EV flow cytometry gating strategy, related to Figure 2 and STAR Methods.** (A) Flow cytometric control samples identifying membrane dye (cell trace blue, CTB) as a detection reagent for EVs. (B) Negative controls showing lack of signal in marker channels of interest. (C) Gating of single and double positive EV populations.

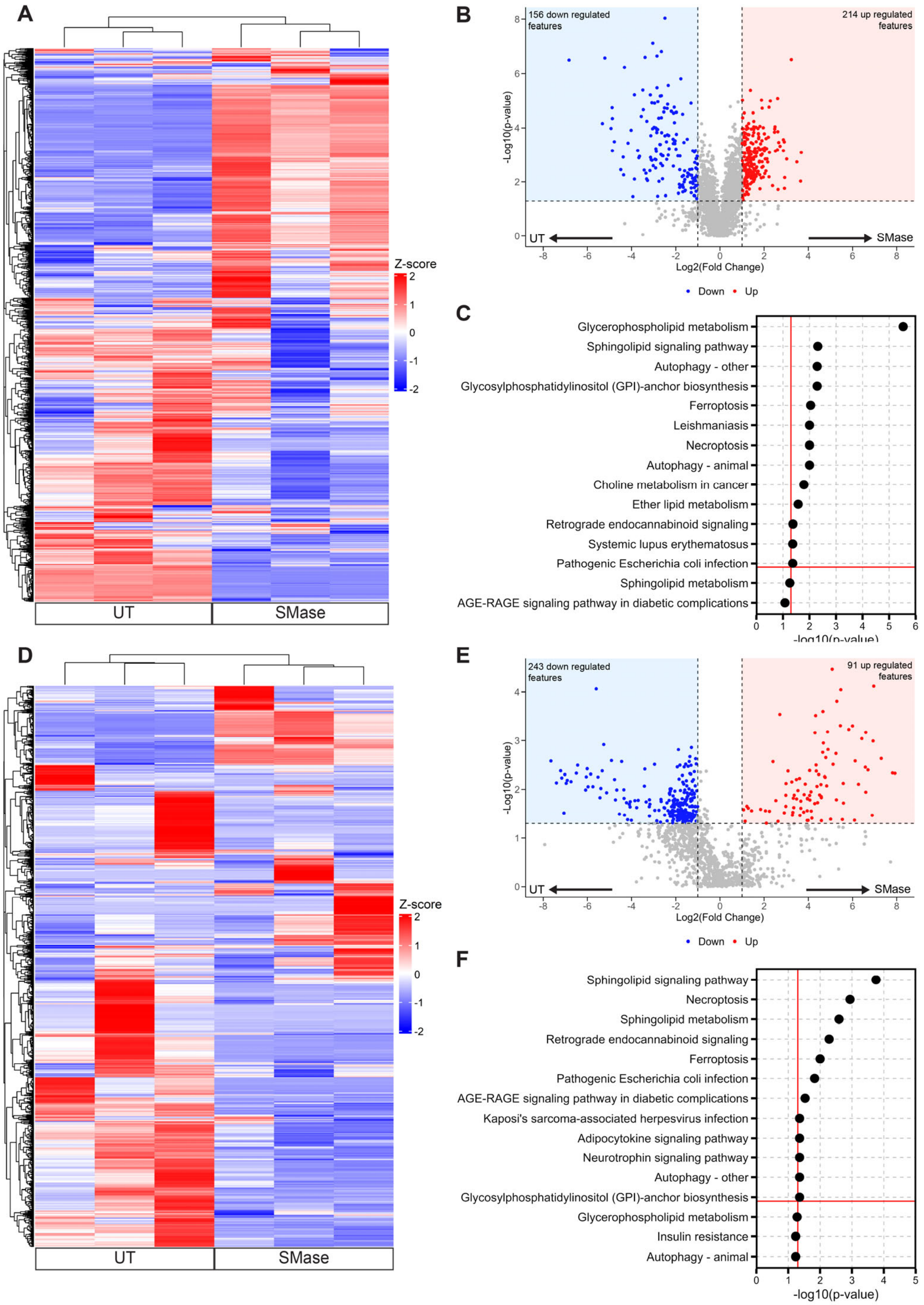


**Figure S3. Comparative lipidomic analysis reveals distinct lipid signatures in SMase-treated MSCs and their derived EVs, related to Figure 3.** (A) Hierarchical clustering heatmap showing differential lipid features in untreated (UT) and SMase-treated MSCs (red: downregulated; blue: upregulated). (B) Volcano plot highlighting significantly altered lipid species in MSCs (red: upregulated; blue: downregulated; gray: unsignificant). (C) Lipid enrichment pathway analysis showing enriched metabolic processes in MSCs. (D) Hierarchical clustering heatmap demonstrating differential lipid profiles between UT and SMase-derived EVs. (E) Volcano plot analysis of EV lipid content showing significantly altered species between treatment groups. (F) Lipid enrichment pathway analysis in EVs. Statistical significance determined by p-values < 0.05; n=3.

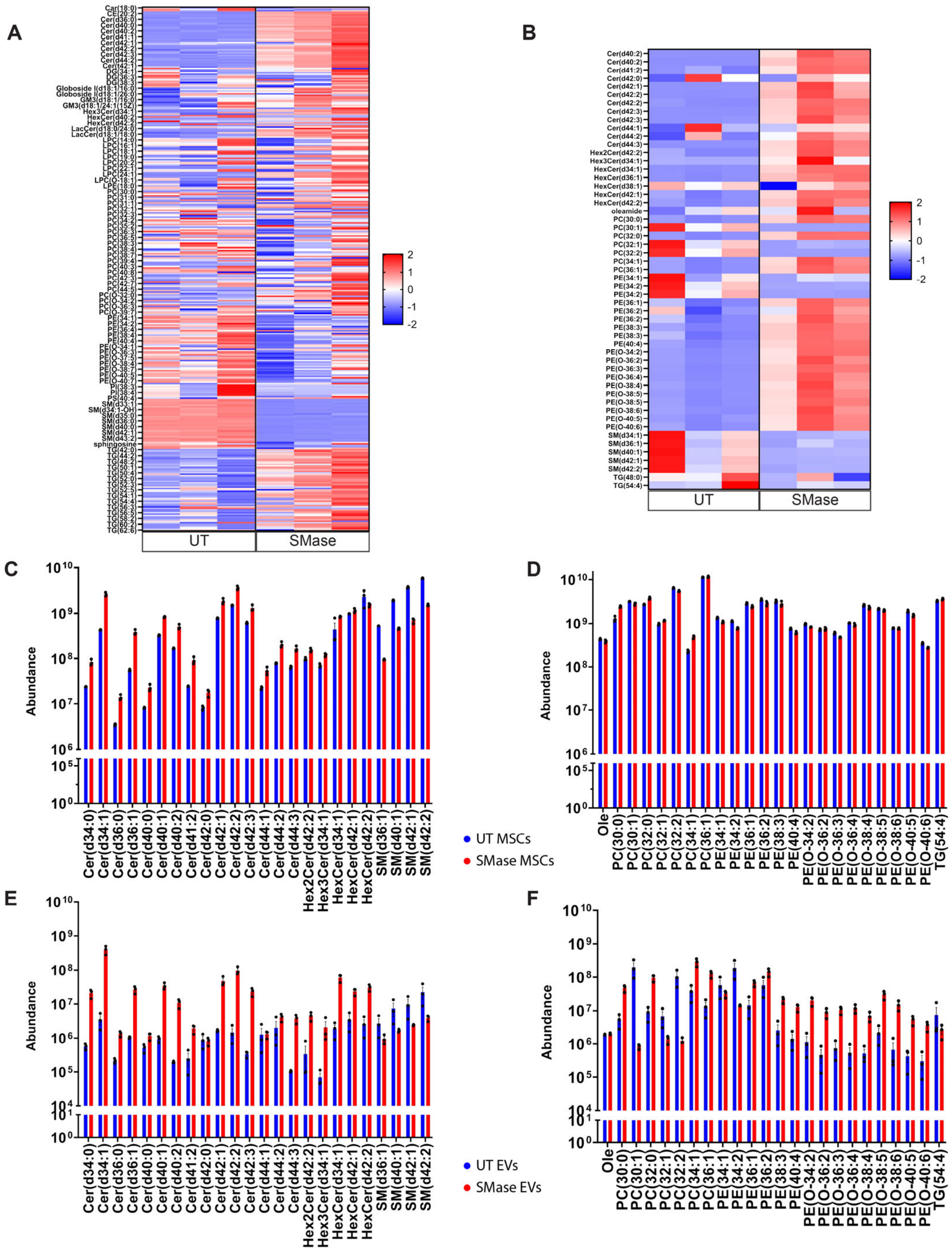


**Figure S4. Annotated lipid profiles shift following SMase stimulation in MSCs and their derived EVs, related to Figure 3.** Z-scored heatmaps of the annotated lipid profiles found in (A) MSCs and (B) MSC-derived EVs. Quantitative analysis of shared annotated sphingolipid and non-sphingolipid abundance values from (C-D) MSCs and (E-F) EVs of both treatments. Ole – oleamide, Car – carnitine, TG – triglyceride, PI – phosphatidylinositol, PE – phosphatidylethanolamine, PE-O – ester linked phosphatidylethanolamine, PC – phosphatidylcholine, PC-O – ester linked phosphatidylcholine, DG – diacylglycerol, LPE – lysophosphatidylethanolamine, LPC – lysophosphatidylcholine, LPC-O – ester linked lysophosphatidylcholine, CE – cholesteryl ester, SM – sphingomyelin, Cer – ceramide, HexCer – hexosylceramide, Sph – sphingosine. n=3.

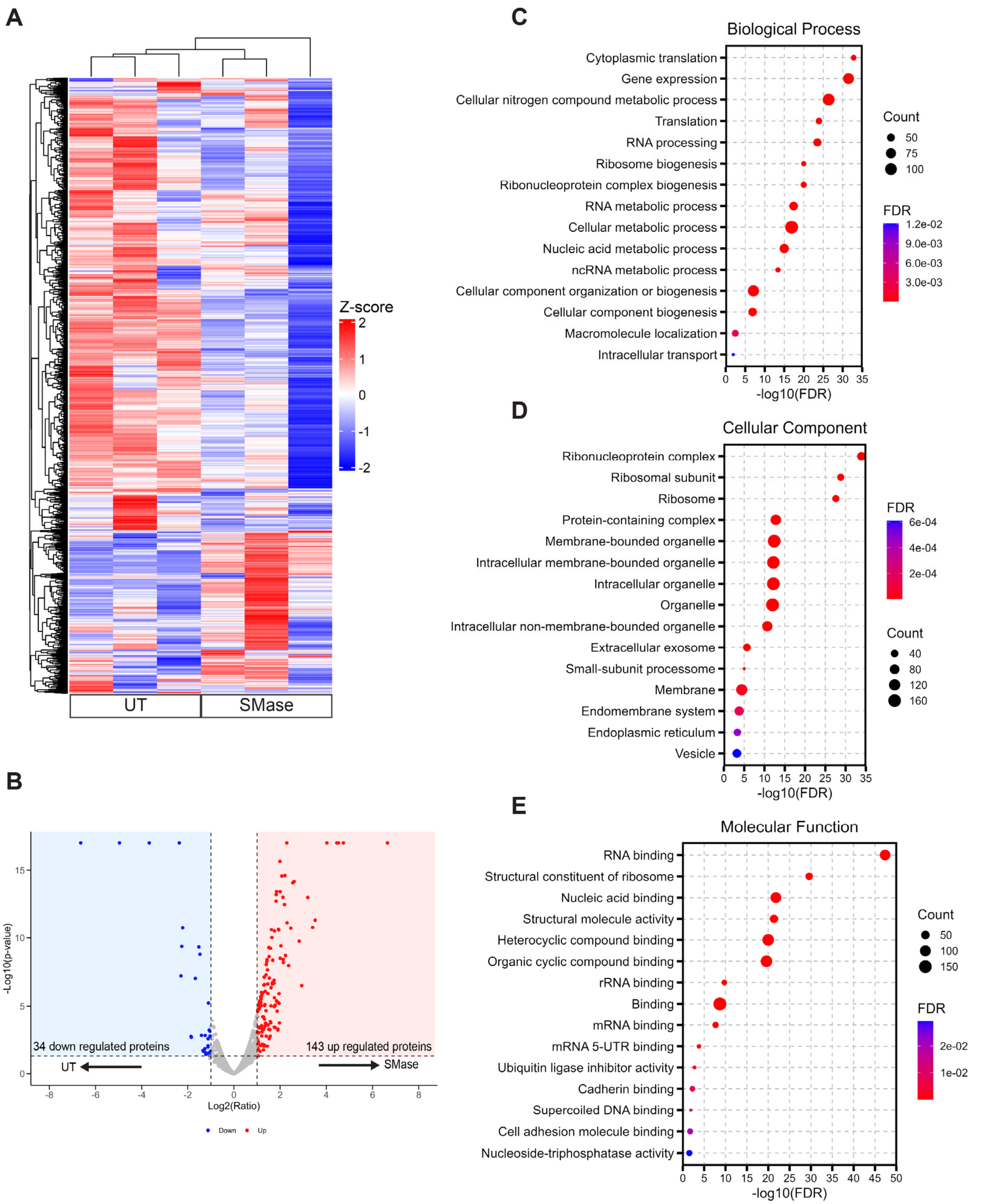


**Figure S5. SMase-treatment induces systematic proteomic remodeling in MSCs with enrichment of RNA processing and membrane organization pathways, related to Figure 4.** (A) Hierarchical clustering heatmap demonstrating differential protein expression patterns between untreated and SMase-treated MSCs (blue: downregulated; red: upregulated). (B) Volcano plot of all detected features (blue: downregulated;

red: upregulated; gray: unsignificant). (C-E) Gene ontology analysis revealing enriched terms in (C) biological processes, (D) cellular components, and (E) molecular functions. Statistical significance determined by false discovery rate-adjusted p-values; dot size indicates gene count; color intensity represents FDR value. n=3.

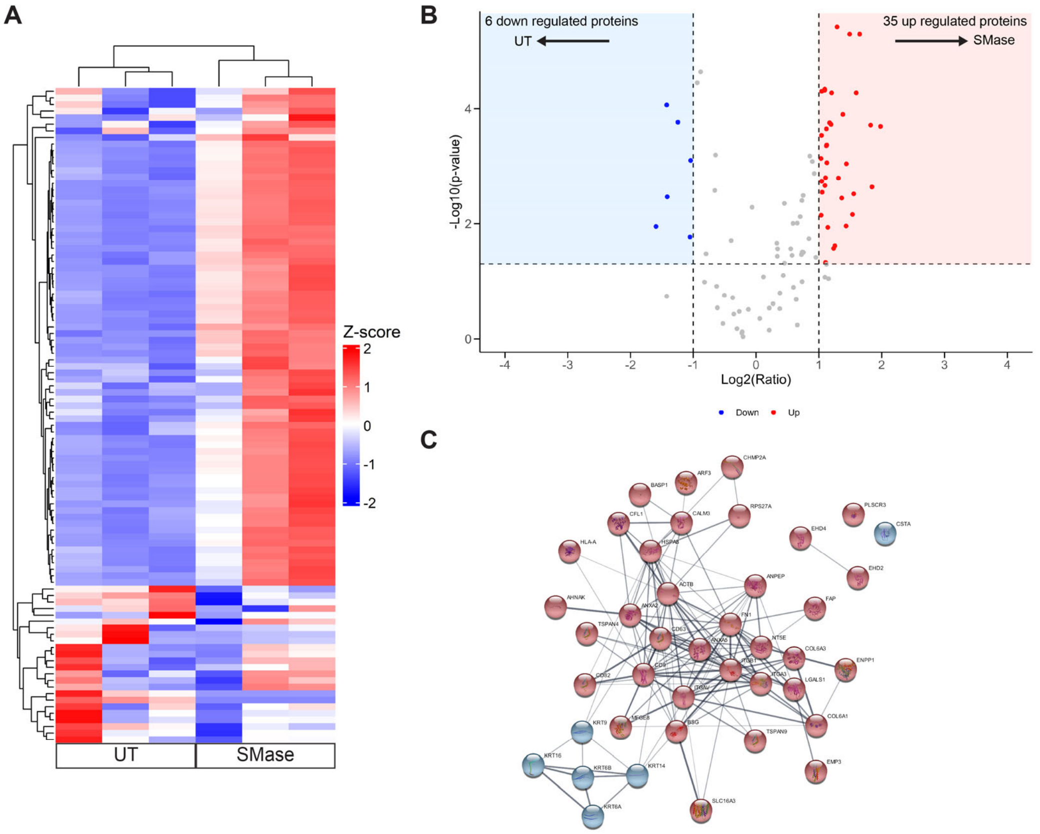


**Figure S6. SMase treatment alters EV protein cargo composition, related to Figure 4.** (A) Hierarchical clustering heatmap showing differential protein expression between untreated and SMase-treated MSC-derived EVs (blue: downregulated; red: upregulated). (B) Volcano plot of all detected features (blue: downregulated; red: upregulated; gray: unsignificant). (C) STRING database generated protein network map depicting the predicted protein-to-protein associations (edges) for proteins up regulated (red) and down regulated (blue) following SMase stimulation. n=3.

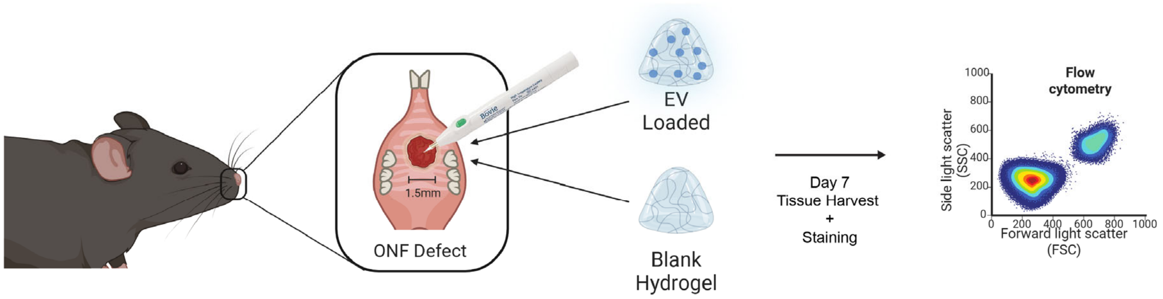


**Figure S7. Local delivery of EV-loaded synthetic hydrogels in a critical murine oral wound model.**

**Table S1. Composition of stable isotope-labeled chemical standards mixture used in UHPLC-MS, related to STAR Methods.**

| **Isotopically labeled lipids** | **CAS number** | **Concentration in stock solution** (µg/ml) |
|---|---|---|
| LPC (18:1(d7)) | 2097561-13-0 | 25 |
| LPE(18:1(d7) | 2260669-47-2 | 5 |
| PC (15:0/18:1(d7)) | 2097561-16-3 | 160 |
| PE (15:0/18:1(d7)) | 2097561-15-2 | 5 |
| PS (15:0/18:1(d7)) | 2260669-40-5 | 10 |
| PG (15:0/18:1(d7)) | 2260669-42-7 | 30 |
| PI (15:0/18:1(d7)) | 2260669-44-9 | 20 |
| CE (18:1(d7)) | 1416275-35-8 | 350 |
| DG (15:0/18:1(d7)) | 2097561-14-1 | 10 |
| TG (15:0/18:1(d7)/15:0) | 2097561-17-4 | 55 |
| SM (18:1(d9)) | 2260669-50-7 | 30 |
| Cholesterol-d7 | 83199-47-7 | 100 |
